# The RNA-binding protein, Imp specifies olfactory navigation circuitry and behavior in *Drosophila*

**DOI:** 10.1101/2023.05.26.542522

**Authors:** Aisha Hamid, Hannah Gattuso, Aysu Nora Caglar, Midhula Pillai, Theresa Steele, Alexa Gonzalez, Katherine Nagel, Mubarak Hussain Syed

## Abstract

Complex behaviors depend on the precise developmental specification of neuronal circuits, but the relationship between genetic programs for neural development, circuit structure, and behavioral output is often unclear. The central complex (CX) is a conserved sensory-motor integration center in insects that governs many higher order behaviors and largely derives from a small number of Type II neural stem cells. Here, we show that Imp, a conserved IGF-II mRNA-binding protein expressed in Type II neural stem cells, specifies components of CX olfactory navigation circuitry. We show: (1) that multiple components of olfactory navigation circuitry arise from Type II neural stem cells and manipulating Imp expression in Type II neural stem cells alters the number and morphology of many of these circuit elements, with the most potent effects on neurons targeting the ventral layers of the fan-shaped body. (2) Imp regulates the specification of Tachykinin expressing ventral fan-shaped body input neurons. (3) Imp in Type II neural stem cells alters the morphology of the CX neuropil structures. (4) Loss of Imp in Type II neural stem cells abolishes upwind orientation to attractive odor while leaving locomotion and odor-evoked regulation of movement intact. Taken together, our work establishes that a single temporally expressed gene can regulate the expression of a complex behavior through the developmental specification of multiple circuit components and provides a first step towards a developmental dissection of the CX and its roles in behavior.

## Introduction

Complex behaviors depend on the development of intricate neural circuits that integrate and store information to generate meaningful behavior. Developmental defects can cause devastating malformations of brain circuitry in humans, with profound consequences for learning and behavior^1, 2^. Conversely, modifications to developmental genetic programs can underlie the evolution of more elaborate brain structures and behaviors from simpler ancestral patterns^3, 4^. However, the relationship between genetic programs for neural development, neural circuit structure, and behavioral capabilities is challenging to study in complex vertebrate brains.

In insects, a conserved brain structure known as the central complex (CX)^5, 6^ has been implicated in many higher-order behaviors, such as locomotion and navigation^6–21^, action selection^22–24^, steering^25, 26^, feeding^27, 28^, courtship^29^, and sleep^30–37^. The CX comprises several conserved structures^38–41^ (Fig. 1A), including the ellipsoid body (EB), which houses a global heading representation^15^, and the protocerebral bridge (PB), which broadcasts this signal to other parts of the CX^42, 43^. The function of the fan-shaped body (FB) is more mysterious, although recent studies have implicated this region in controlling locomotion^25^, computing allocentric variables^20, 21^, and specifying behavioral goals^44, 45^. A scaffold of columnar neurons links the different regions of the CX. Columnar inputs to the fan-shaped body encode both heading information derived from the protocerebral bridge and optic flow or airflow information from the paired noduli (NO)^20, 46^. The FB also receives many tangential inputs that carry information from various parts of the dorsal brain and encode non-spatial stimuli such as odors^45^, tastants^28^, and internal states^32^. Local fan-shaped body neurons known as hΔ cells integrate input from columnar and tangential inputs^45, 47^. Patterned stimulation of a subset of local neurons can drive reproducible navigation in an allocentric direction, implying that these neurons may encode the animal’s goals^44, 45^. Transmission electron microscopy (TEM) reconstruction of the hemibrain connectome^48^ has identified 224 morphological and 262 connectivity types in the CX^47^, many of which are primarily conserved in other insects such as bees^49^. The near-crystalline structure of the CX, along with its role in complex behaviors, make it an ideal model for investigating the relationship between neural development, circuit structure, and behavior. Clonal analysis of adult Drosophila brain has shown that CX is derived from many lineages, and many clones innervate same neuropils giving rise to densely interconnected neuropils^50, 51^. In addition, lineage analysis has shown that the Drosophila brain is composed of anatomically distinct compartments with lineage boundaries and compartment boundaries defined by dense layers of glial cells^52, 53^. However, how the developmental programs govern formation of a functional circuit has not been established.

**Figure1.**
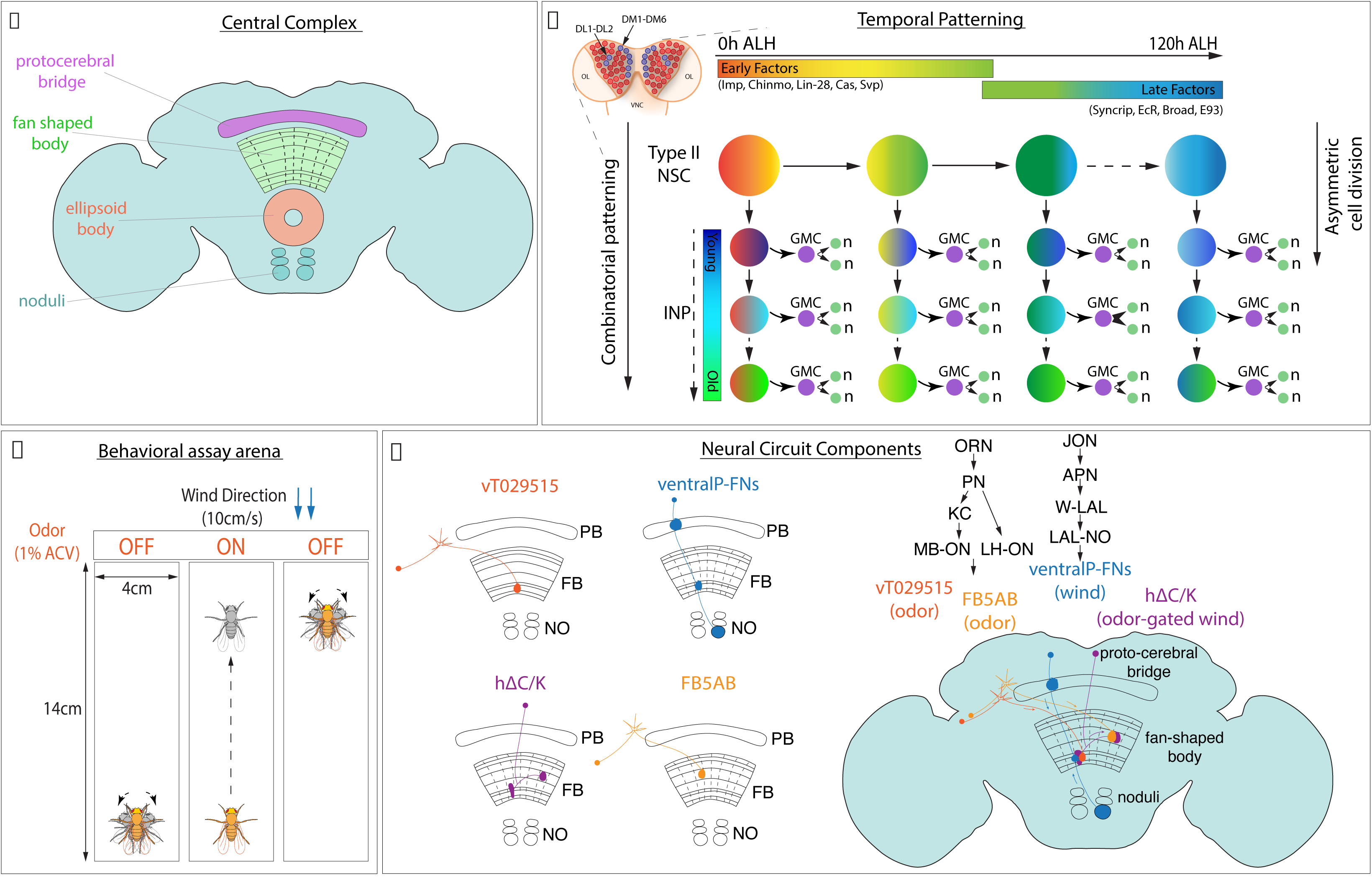
Schematics depicting central complex, division pattern, and temporal patterning in Type II neural stem cells (NSCs), olfactory navigation behavioral assay arena, and circuit components: **A.** *Drosophila* adult brain central complex with labelled neuropils, protocerebral bridge (magenta), fan shaped body (green), ellipsoid body (orange), and the paired noduli (blue). **B.** Type II NSCs in the larval brain (blue, 8 per lobe: DM1-6, and DL1-2) divide asymmetrically, producing another NSC and an intermediate neural progenitor (INP). INP divides and gives rise to another INP and a ganglion mother cell (GMC), which eventually forms two neurons and/or glia. Type II NSCs express early and late factors in a time-dependent manner starting from 0h ALH to 120h ALH referred to as early and late factors, respectively. In addition to temporal patterning in NSCs, INPs also express different factors in a birth order-dependent manner along with NSC factors and give rise to a combinatorial code. **C.** Schematic of the behavioral assay arena; walking flies are observed in a rectangular chamber (14cmX4cm) with the constant wind flowing from one direction at 10cm/s, and odor is provided in 10s pulse (1% ACV) centered in a 70s trial. The flies tend to walk upwind towards the odor source when the odor cue is ON and demonstrate exploratory behavior by turning right or left when the odor cue is OFF^79^. **D.** CX neurons of the olfactory navigation circuit investigated in this study. These neurons innervate many neuropils of the CX. Long-field FB input neurons encode odor, ventral P-FNs encode wind, and the common downstream neurons are the local hΔC/K neurons^45^

Most adult *Drosophila* CX neurons arise from 8 bilateral larval neural stem cells known as Type II neural stem cells (based on their location: dorsomedial, DM1-6, and dorsolateral, DL1-2). In contrast to the more common Type I neural stem cells, which divide to self-renew and generate a pair of differentiated neural progeny, Type II neural stem cells divide asymmetrically to self-renew and produce an intermediate neural progenitor (INP)^54, 55^. INPs further divide multiple times to generate 8-12 progeny and have been shown to express different factors in a birth order dependent manner and play a role in generating neural identity^56–64^ (Fig. 1B). The dual division pattern of Type II neural stem cells is shared by the outer radial glia (oRG), stem cells that generate the primate and human cortex^65–67^. And the intermediate progenitors have been reported in human cortex as well^68–70^. Type II-like neural stem cells are also found in grasshoppers and beetles^71–74^, making them evolutionarily-conserved progenitors that generate higher-order brain centers. Type II neural stem cells produce a significant proportion of CX lineages^75^. The CX columnar neurons arise from four bilateral Type II neural stem cells called DM1-4, while tangential neurons arise from mixed lineages, with a significant contribution from DL1^53, 75^.

To generate diverse classes of neuronal types over time, Type II neural stem cells express sets of temporally regulated transcription factors and RNA binding proteins (RBPs)^76, 77^. Using RNA-Seq methods, we and others have identified about a dozen temporally expressed genes in Type II neural stem cells^76, 77^. These factors can be divided into two groups: early factors detected from 0-60h ALH (after larval hatching) and late factors from 60-120h ALH (Fig. 1B). Early expressed factors include the RBPs Imp and Lin-28, and the transcription factors Chinmo, Seven-up, and Castor. Late factors include the RBP Syncrip and the transcription factors, Ecdysone receptor (EcR), Broad, and Ecdysone-induced protein 93 (E93)^76^. In addition, extrinsic hormonal signaling via timely expression of the receptor EcR at ∼55h ALH induces an early-to-late gene transition in Type II neural stem cells^76^. Opposing gradients of Imp and Syncrip expression have previously been shown to regulate the number and morphology of neurons in the CX^77^. However, how these temporally expressed factors contribute to functional circuit formation, and behavior is unknown.

Here we use an innate goal-oriented behavior— olfactory navigation— and an associated neural circuit to gain insight into how developmental programs specify neural types and shape circuit structure to determine behavioral output. When a hungry fly encounters an attractive odor, it will turn and increase its velocity upwind^78, 79^, a conserved behavior observed across many arthropods^80^ (Fig. 1C). In a previous study, we implicated several CX neuron types in this behavior^45^ (Fig. 1D, Table 1). The sensory cues required for this behavior— odor and wind direction— are encoded by different types of FB inputs. Tangential input neurons (FB5AB, 65C03, 12D12, VT029515) encode odor^45^, while columnar input neurons (ventral P-FNs) encode wind direction^46^. A set of local FB neurons labeled by the line VT062617-GAL4 integrates these cues and is required for persistent upwind tracking in flies^45^. In that study, we identified this line as labeling hΔC neurons; however, single-cell clones revealed that this line also labels hΔK neurons (Supp. Fig. 1). We, therefore, refer to this population as hΔC/K. Here we show that all elements of this circuit are derived from Type II neural stem cells and that Imp is required for the specification of both tangential and columnar neurons targeting the ventral fan-shaped body. We find that different circuit elements show distinct requirements for Imp either in neural stem cells or post-mitotically. Imp knock-down in Type II neural stem cells shapes the gross morphology of the CX and the expression of peptides within specific layers of the fan-shaped body. Finally, we show that manipulating Imp expression in Type II neural stem cells profoundly impairs olfactory navigation behavior. Collectively, our findings demonstrate a novel role of Imp in the formation and function of an olfactory navigation circuit and trace the developmental origins of distinct components of a behavioral circuit.

**Table 1:**

| **GAL4/LexA line** | **Neuron class labeled** |
| --- | --- |
| VT029515 | Ventral FB tangential inputs |
| VT062617 | hΔC/K FB local neurons |
| 21D07 | FB5AB (a dorsal FB tangential input) |
| 15E12/30E10 | Ventral P-FNs, FB column inputs |
| 65C03 | Dorsal FB tangential inputs |
| 12D12 | Dorsal and ventral FB tangential inputs |

## Results

### Type II neural stem cells generate multiple components of an olfactory navigation circuit

The majority of the neurons populating the adult CX are derived from 16 larval Type II neural stem cells^75^ (schematics shown in Fig. 2A). We first asked whether previously identified elements of an olfactory navigation circuit in the CX arise from these stem cells and determined their lineage. We used an intersectional genetic strategy to express GFP in specific neuron classes in a Type II specific manner (genetic scheme shown in Fig. 2B). Wor-GAL4 is expressed in all neural stem cells, while Ase-GAL80, the repressor for GAL4, is expressed only in Type I neural stem cells, leaving GAL4 active in only Type II neural stem cells. Flippase now expressed only in Type II neural stem cells permanently flips out a stop sequence between LexAop and GFP, making Type II progeny eligible to express GFP under LexA control. A neuron class-specific LexA can then drive expression of GFP only if those neurons are derived from Type II neural stem cells.

**Figure 2.**
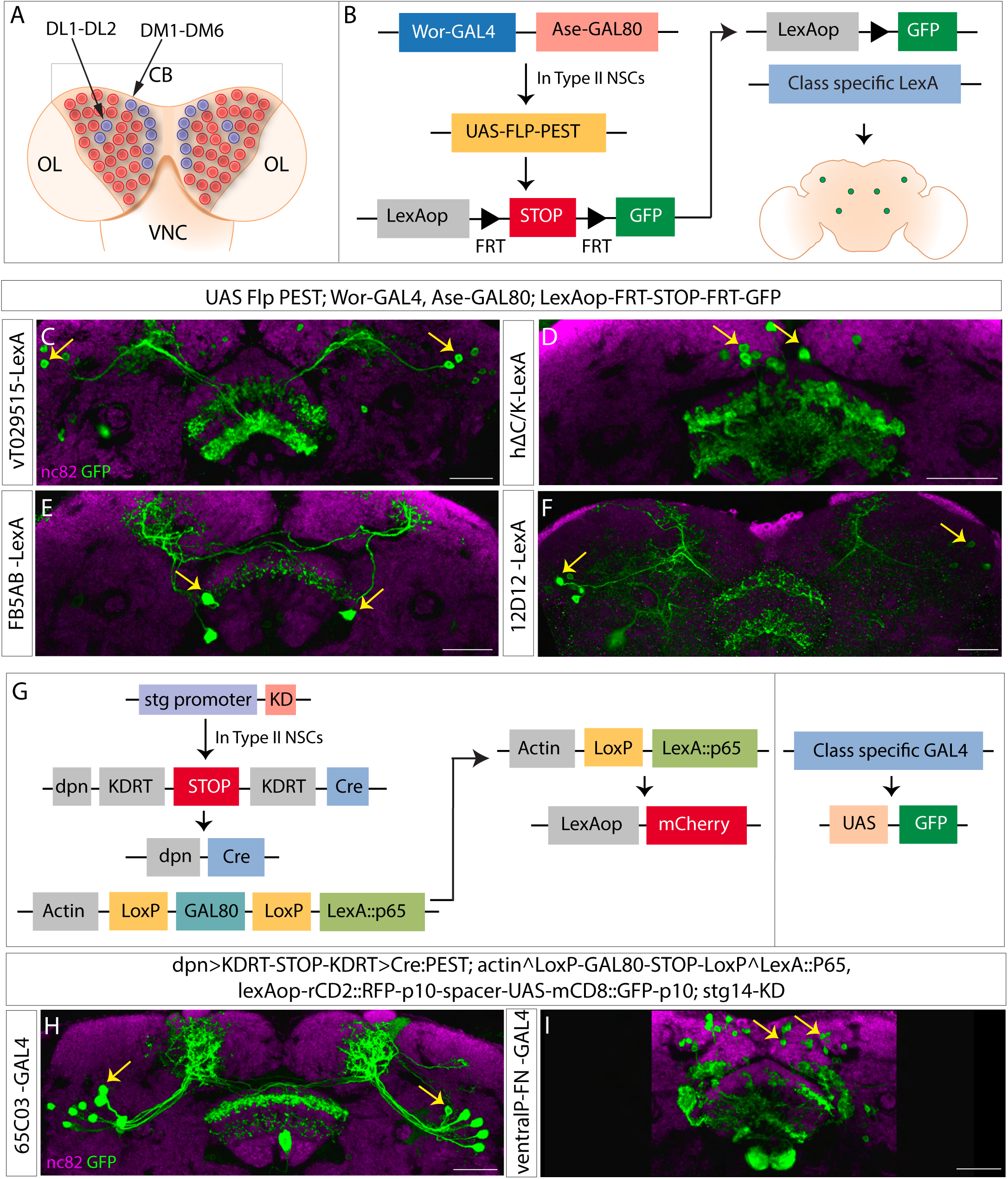
Lineage analysis of olfactory navigation circuit components: **A.** Schematics showing 8 Type II NSCs per larval brain lobe (blue) DM1-6 and DL1-2, among other NSCs (Type 0 and Type I) in red. **B.** Genetic scheme for Type II lineage analysis using Type II specific Flp-out. Ase-GAL80 active in Type I NSCs leaves Wor-GAL4 active only in Type II NSCs, Flippase downstream of UAS sequence is activated in Type II specific manner and removes the STOP sequence between two FRT sites and expresses reporter GFP in class-specific manner when crossed to a class-specific LexA driver line. **C-F.** Confocal images showing neuron types expressing reporter GFP in Type II dependent manner. **G.** Genetic scheme for Type II lineage filtering using CLIn technique. Type II specific promoter fragment combined with KD recombinase removes STOP sequence by a flip event and leaves Cre recombinase active in Type II NSCs. Cre generates Type II NSC clones positive for LexA::p65.GAL80 is active in the rest of the cells, and GAL4 expresses GFP as a reporter in class specific manner. We did not stain for mCherry here. **H-I.** Confocal images showing neuron types expressing reporter GFP in Type II dependent manner. nc82 stain in magenta counterstains the brain, a stack of 2 slices was taken for visualizing the neuropil with nc82. GFP reporter is shown in green. Cell bodies are indicated by yellow arrows. The genotypes are shown at the top, and the abbreviated genotypes for neuron class are shown at the left. Scale bars correspond to 30μm.

Using this strategy, we found that several components of CX olfactory navigation circuitry are derived from the Type II neural stem cells. These include multiple tangential FB inputs that encode odor (VT029515, FB5AB, 12D12), as well as local neurons that integrate odor and wind information (hΔC/K, Fig. 2C-F). We used a different genetic strategy for two additional neuron classes to determine their lineage, as we had only GAL4 and not LexA lines for these classes at the time. This strategy is based on the Cell class-lineage intersection (CLIn) transgenic system^81^. Here multiple flip-out events mediated by Flippase, Cre, and KD recombinases restrict the clonal analysis to Type II neural stem cells. The Type II specific promoter Stg drives the expression of the reporter A-mCherry in a lineage-specific manner, and a class-specific GAL4 drives the expression of reporter B-GFP in Type II dependent manner (genetic scheme shown in Fig. 2G). This analysis showed that two additional components of olfactory navigation circuitry, an odor-encoding dorsal tangential input (65C03) and a wind-encoding columnar input (ventral P-FNs), also arise from Type II neural stem cells (Fig. 2H and I). Taken together, our lineage analysis shows that multiple elements of an olfactory navigation circuit in the CX arise from Type II neural stem cells.

### Mid-aged dorsolateral Type II neural stem cells produce odor-encoding ventral FB tangential input neurons

Our lineage analysis revealed that several tangential FB input neurons are derived from Type II neural stem cells. We decided to examine the lineage and birth time of one population of ventral tangential FB inputs (VT029515) more closely. To do this, we combined CLIn with a heat-sensitive Flippase, allowing us to determine the birth time of a specific neuronal class (complete genetic scheme shown in Fig. 3A). Heat shock was given by shifting larvae to 37°C at different time points (Fig. 3B).

**Figure 3.**
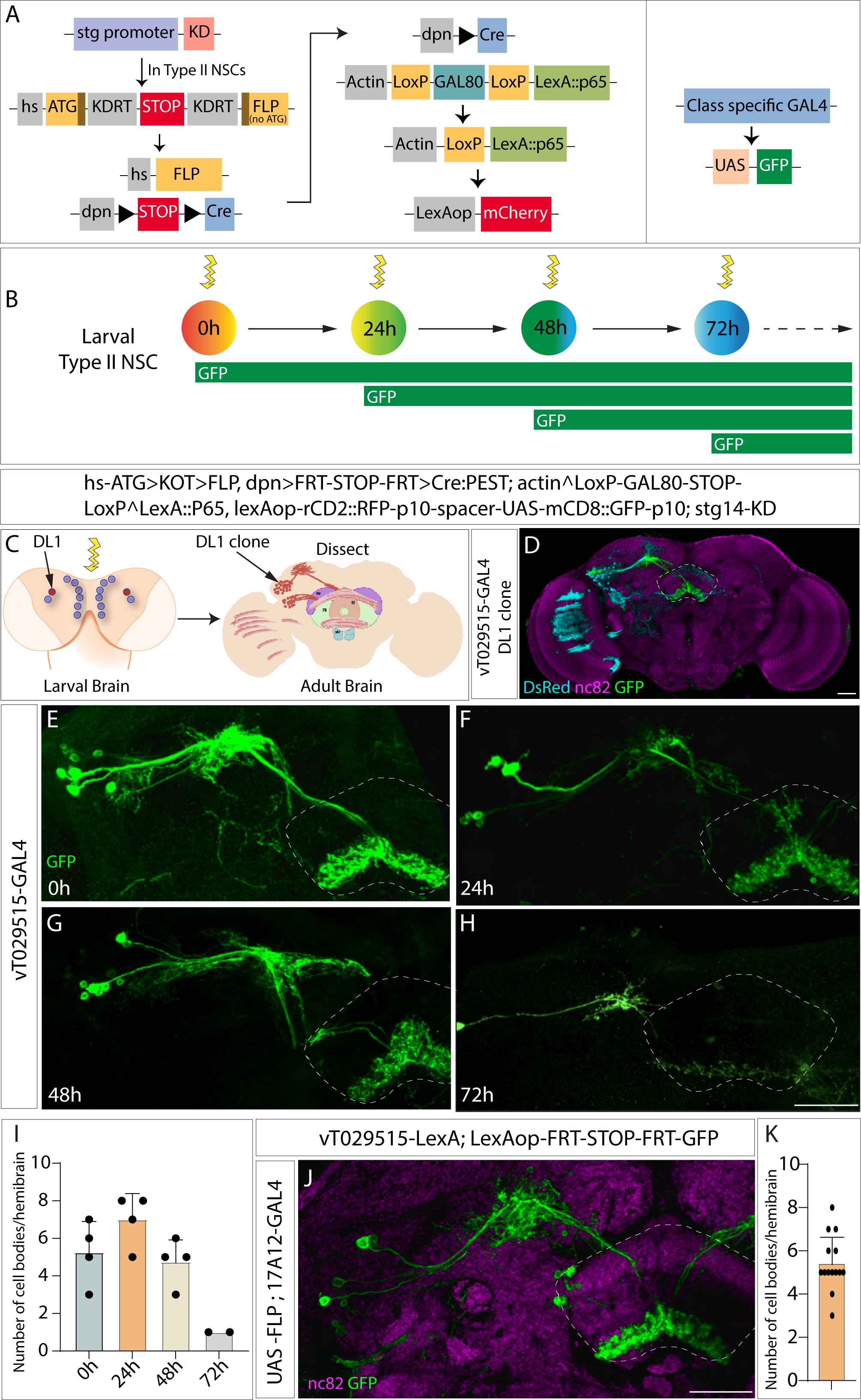
Birth-dating vT029515 long field tangential input neurons using cell-class lineage intersection system: **A.** Schematics describing the genetics of the CLIn system combined with a heat shock promoter for genetic birth-dating: Type II specific stg promoter fragment combined with KD recombinase removes STOP sequence by a flip event, and an intact Flippase is reconstituted, this combined with heat-sensitive promoter removes STOP sequence and leaves Cre-recombinase active in Type II NSCs in a heat shock dependent manner. Cre in turn generates Type II NSC clones positive for LexA::p65. GAL80 is active in the rest of the cells, and GAL4 expresses GFP as a reporter in class-specific manner, while as a reporter mCherry downstream of LexAop sequence is expressed in a lineage-specific manner. **B.** Schematics representing heat shock given at different time points (ALH-after larval hatching), yellow arrow indicates heat shock, and GFP expression window is shown in green. **C.** Schematics showing the map of the progeny of the DL1 clone populating dorsal and ventral layers of the FB^75^ in the adult brain. **D.** 0h DL1 clone of CLIn crossed to VT029515 GAL4 labeling vT029515 neurons, nc82 counterstains the brain, a stack of 2 slices was taken for visualizing the neuropil with nc82, DsRed in cyan labels the DL1 progeny, GFP in green labels vT029515 neurons. **E-H.** 0h, 24h, 48h, and 72h ALH clones, GFP in green labels vT029515 neurons. **I.** Quantification of number of cell bodies counted per hemibrain at indicated time points (n=data points indicated on the graph). **J.**17A12-GAL4 flipped vT029515 neurons, nc82 counterstains the brain, a stack of 2 slices was taken for visualizing the neuropil with nc82. GFP in green labels vT029515 neurons. **K.** Quantification of number of vT029515 neurons per hemibrain flipped by 17A12-GAL4. The genotypes are shown at the top, and the abbreviated genotypes for neuron class are shown at the left. Scale bars correspond to 30μm.

Single CLIn clonal analysis revealed that most VT029515 neurons are derived from the DL1 neural stem cell (number of cell bodies=7) (Fig. 3D, Video1), and a few are derived from DMs (number of cell bodies= 2, data not shown). The progeny of DL1 neural stem cells have been earlier reported to innervate the FB in two distinct bundles, dorsal and ventral^75^ (schematics shown in Fig. 3C). Next, we performed genetic birth-dating by inducing flip-out events via temporal heat shock at different developmental time points in larvae: 0h, 24h, 48h, and 72h ALH. This genetic birth-dating revealed that most Type II derived VT029515 neurons are generated between 48h-72h ALH (Fig. 3E-I), an interval spanning the early-to-late gene expression switch at 60h in all Type II neural stem cells^76^. Combining our clonal analysis and genetic birth-dating assays, we conclude that vT029515 input neuron types are born from mid-aged DL1 neural stem cells. To confirm this result, we used an intersectional genetic strategy. The line 17A12-GAL4 expresses selectively in the DL1 and DL2 stem cells during development (unpublished). We used this line to express Flippase and excise an intervening STOP cassette from LexAop-FRT-STOP-FRT-GFP only in DL1 and DL2 neural stem cells. We combined these transgenes with VT029515-LexA to label ventral FB tangential input neurons. Upon expression of Flippase by 17A12-GAL4, we observed 4-8 vT029515 neurons labelled (n=14 hemibrains) (Fig. 3J-K). Variation in the number of neurons labeled might arise from variability in Flippase expression. These findings support our hypothesis that most vT029515 neuron types are born from DL1 Type II neural stem cells, while some are derived from other Type II neural stem cells.

### Imp regulates the specification of odor-encoding ventral FB tangential input neurons

Early Type II neural stem cells express the transcription factors Chinmo, Castor, Seven up, and the conserved RBPs, Imp and Lin-28. To test our hypothesis that an early neural stem cell factor regulates the fate of VT029515 neurons, we focused on a conserved RBP, IGF-II mRNA-binding protein, Imp^82^. We asked if Imp expression during development determines the number and morphology of VT029515 neurons. We expressed either UAS-ImpRNAi (2 copies) or UAS-Imp in Type II neural stem cells throughout development using the driver Pointed-GAL4, and assayed animals at adult stages. We combined this manipulation with a LexA-based GFP reporter to examine the number and morphology of VT029515 neurons (VT029515-LexA). We found that Imp manipulations profoundly affected the specification of VT029515 neurons. In control animals, we counted about 6-7 cell bodies per hemisphere (Fig. 4A, A’). These neurons were completely absent in Type II>ImpRNAi flies (Fig. 4B, B’). To test if Imp is sufficient for generating VT029515 neurons, we ectopically expressed Imp in Type II neural stem cells, Type II>UAS-Imp. In these animals, Imp is now expressed in young as well as late Type II neural stem cells^83^. We observed a ∼3-fold increase in the number of VT029515 cell bodies with Imp overexpression. These ectopic neurons formed part of the same axon bundle and similarly innervated the ventral layer of the fan-shaped body (Fig. 4C, C’).

**Figure 4.**
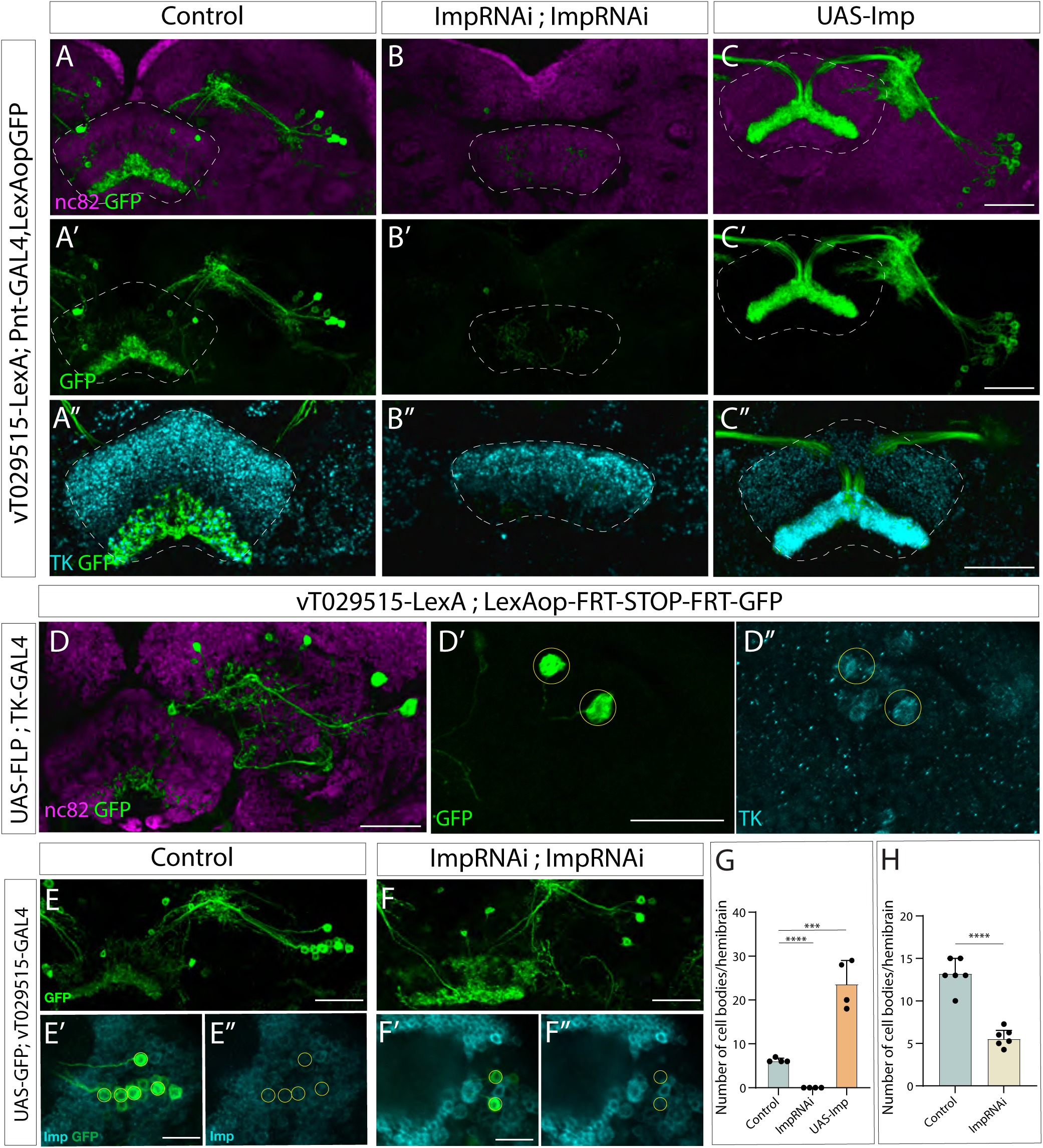
Imp is required for specifying and maintaining the identity of odor encoding vT029515 long field input neurons: **A-C’**. vT029515 neurons labelled by reporter GFP in control **(A)**, Imp knock-down **(B),** and Imp overexpression **(C)**, nc82 counterstains the brain, and GFP is shown in green. **A’’-C’’.** vT029515 neuron arbors shown by GFP expression in green, neuropeptide TK staining shown in cyan colocalizing with GFP for control **(A’’)**, Imp knock-down, expression observed in dorsal layers of FB **(B’’)** and Imp overexpression, TK expressed in thick bundle colocalizing with GFP expression **(C’’)** **D.** vT029515 neurons labelled with GFP via conditional expression of GFP with 17A12-GAL4, nc82 counterstains the brain. Zoomed in view of vFB cell bodies, GFP in green **(D’),** and TK expression **(D’’)** **E-F’’.** vT029515 neurons labelled by reporter GFP are shown in green, Imp expression is shown in cyan in control **(E-E’’)** and Imp knock-down **(F-F’’)**. For E’-E’’ and F’-F’’, the Scale bar corresponds to 15μm. **G-H.** Quantification of number of cell bodies per hemibrain for Type II specific knock-down **(G)** and post-mitotic knock-down **(H)**. n=data points indicated on the graphs. (Students t-test). The abbreviated genotypes are shown at the left and top, respectively. Scale bars for the rest correspond to 30μm. The dashed outline shows FB in A-C’’ and cell bodies in D’-D’’ and E’-F’’.

To determine whether the ectopic and endogenous VT029515 neurons have the same identity, we stained for a neuropeptide Tachykinin (TK), known to be expressed in the ventral layers of the FB^84^. We observed that the VT029515 arborization in the ventral layers of the FB colocalizes with TK immunofluorescence (Fig. 4A’). Interestingly, TK expression was lost in the ventral layers upon Imp knock-down, suggesting an essential role of Imp in specifying neurons with TK neuropeptide identity (Fig. 4B’’). In contrast, Imp gain of function increased TK staining in the ventral layers, which co-localized with the innervations of the ectopic VT029515 neurons (Fig. 4C’’). To confirm that VT029515 neurons express TK, we drove Flippase with TK-GAL4 and used vFB-LexA>LexAop-FRT-stop-FRT-GFP to express GFP only in VT029515 neurons that also express TK-GAL4. We observed 1-3 GFP-positive neurons per hemibrain (n=6), indicating that VT029515 are indeed TK positive (Fig. 4D-D’’). Together these results suggest that ectopic VT029515 neurons are morphologically and molecularly similar to normal VT029515 neurons, and Imp likely regulates TK neuropeptide expression in the FB. (Fig. 4A-C’’ and G).

We observed that VT029515 neurons continue to express low levels of Imp in adulthood (Fig. 4E-E’’). This led us to ask if Imp is required post-mitotically in these cells to maintain their function. To address this question, we knocked down Imp post mitotically in these neurons using VT029515-GAL4. We observed a decrease in cell body number in Imp knock-down compared to control, and Imp is undetectable in the small number of surviving neurons. (Fig. 4F-F’’ and H). We conclude that Imp is also required post-mitotically to maintain the identity of VT029515 neurons.

### Distinct roles of Imp in regulating ventral and dorsal circuit elements

We next asked whether Imp also specifies other essential neuron types of olfactory navigation circuitry. Using a similar strategy to the one described above, we knocked down or overexpressed Imp during development using Pointed-GAL4, a broad driver for Type II neural stem cells, and assayed effects on each circuit element using LexA drivers and a GFP reporter in adult brains. Interestingly, we found that Imp knock-down in Type II neural stem cells caused a loss of wind-sensitive columnar neurons known as ventral P-FNs (here labeled by 30E10-LexA). However, unlike VT029515 neurons, the number of cell bodies did not change upon Imp overexpression (Fig. 5A-C).

**Figure 5.**
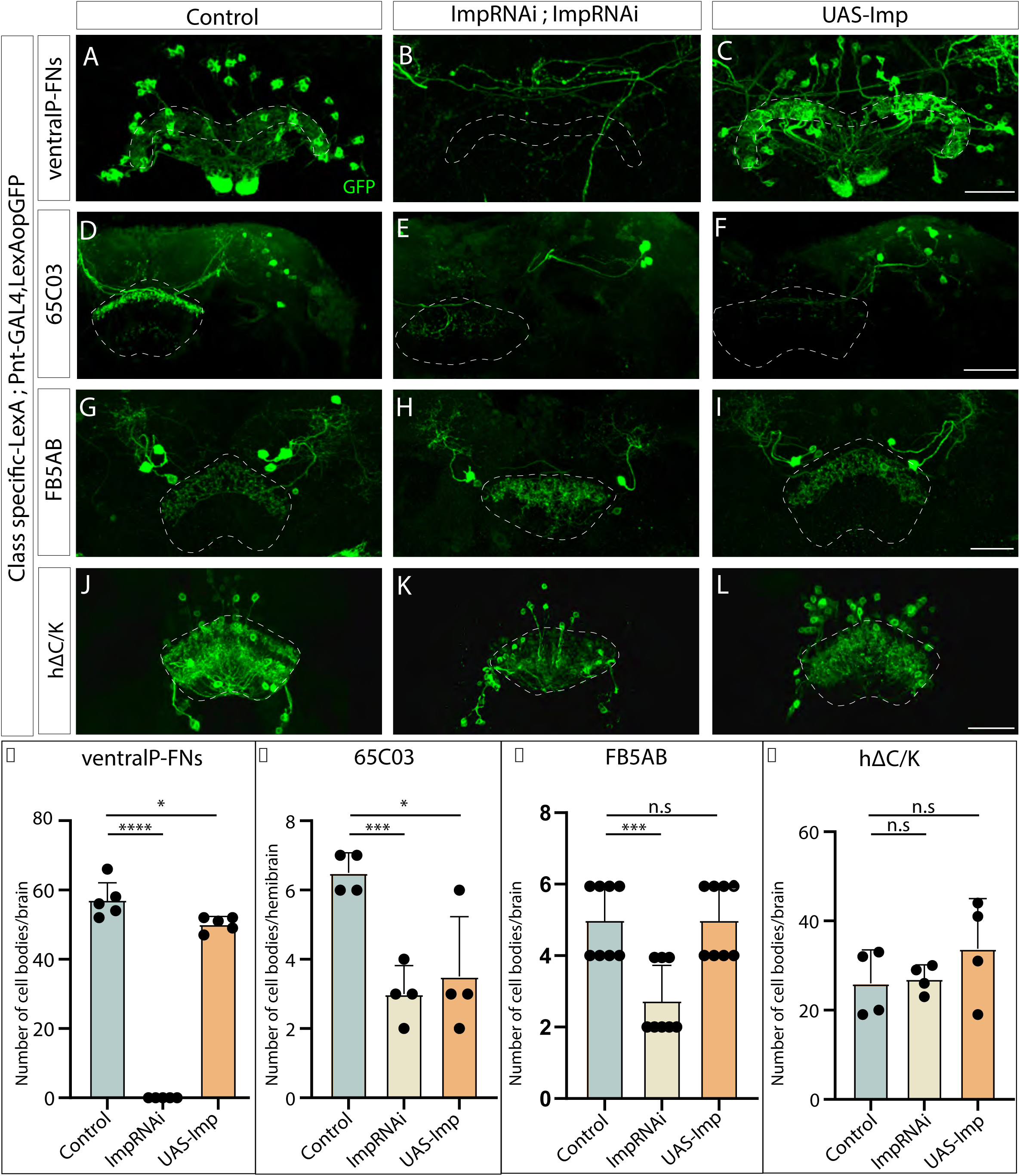
Imp regulates specification and morphology of other neural types in the olfactory navigation circuit: **A-C**. Ventral P-FNs ∼40 neurons labelled by reporter GFP shown in control **(A)**, no neurons labelled in Imp knock-down **(B),** and ∼40 neurons labelled in Imp overexpression **(C)** **D-F.** 65C03 ∼7 neurons labelled by reporter GFP shown in control **(D)**, ∼4 neurons labelled in Imp knock-down with defective morphology of the neurites in FB **(E),** and ∼4 neurons labelled in Imp overexpression **(F)** **G-I.** FB5AB dorsal FB input neurons 4-6 neurons labelled by reporter GFP in control **(G)**, 2-4 neurons labelled in Imp knock-down, with the neurites arborizing FB defectively **(H),** and 4-6 neurons labelled in Imp overexpression **(I)** **J-L.** hΔC/K neurons labelled by reporter GFP in control (J), neurites arborize multiple layers of FB with no clear demarcation in Imp knock-down **(K)** and Imp overexpression **(L)**. GFP is shown in green, the abbreviated genotypes are shown at left. Scale bars correspond to 30μm. **M-P.** Quantification of number of cell bodies. Abbreviated neuron names are indicated at the top. n=data points indicated on the graphs. (Students t-test).

Both dorsal and ventral FB tangential input neurons respond to odor; next, we wanted to know whether Imp regulates the fate of both dorsal and ventral tangential inputs. We first focused on 65C03 neurons, which are 6-8 in number per hemibrain, and project strongly to the dorsal layers and weakly to the ventral layers of the FB. We found that both Imp knock-down and overexpression decreased the number of tangential inputs labeled by 65C03-LexA (Fig. 5D-F). In Imp knock-down flies, we observed only a single projection layer. Moving to the FB5AB dorsal input neurons, we found that 21D07-LexA labels 4-6 neurons projecting to dorsal FB per brain. We observed 2-4 cell bodies in Imp knock-down flies; this could be a decrease in FB5AB neurons or loss of another subpopulation labelled by this LexA line. In Imp knock-down flies, the neurites of these neurons had severely impaired morphology. Imp overexpression did not alter the number of FB5AB tangential inputs labeled by 21D07-LexA, and the morphology was unaffected (Fig. 5G-I). The number of local hΔC/K neurons labeled by VT062617-LexA did not change upon Imp knock-down or overexpression (Fig. 5J-L). However, the morphology of hΔC/K neurons was altered. We observed that changes in Imp levels caused axonal projections of hΔC/K to span multiple FB layers, and the division of dendrites and axons into ventral and dorsal layers was lost. (Fig. 5K) Together, these data suggest that Imp expression during development has the most potent effects on neurons innervating the ventral layers of the fan-shaped body, including both tangential inputs (VT029515) and columnar inputs (ventral P-FNs). Manipulating Imp expression had distinct effects on each neuron type, suggesting that Imp plays multiple roles and acts via multiple mechanisms to regulate cell fate and morphology. These defects suggest that neurons still present in Imp loss-of-function may have altered connectivity.

### Imp regulates CX neuropil morphology and neuropeptide distribution

Our previous results show that Imp loss-of-function influences the morphology of the neurite projections for many of the elements within a CX olfactory navigation circuit. To test if the adult CX is more broadly altered by Imp knock-down in Type II neural stem cells, we compared neuropil (nc82) stains in knock-down and control flies. We found that Imp knock-down results in severely defective CX morphology. The PB is fragmented, the EB is only partially formed, the NO is almost entirely absent, and the FB is smaller, with no clear demarcation of layers (Fig. 6A-I). The FB phenotype is reminiscent of cortical layer deformations observed in Imp1 knock-out mice^85^.

**Figure 6:**
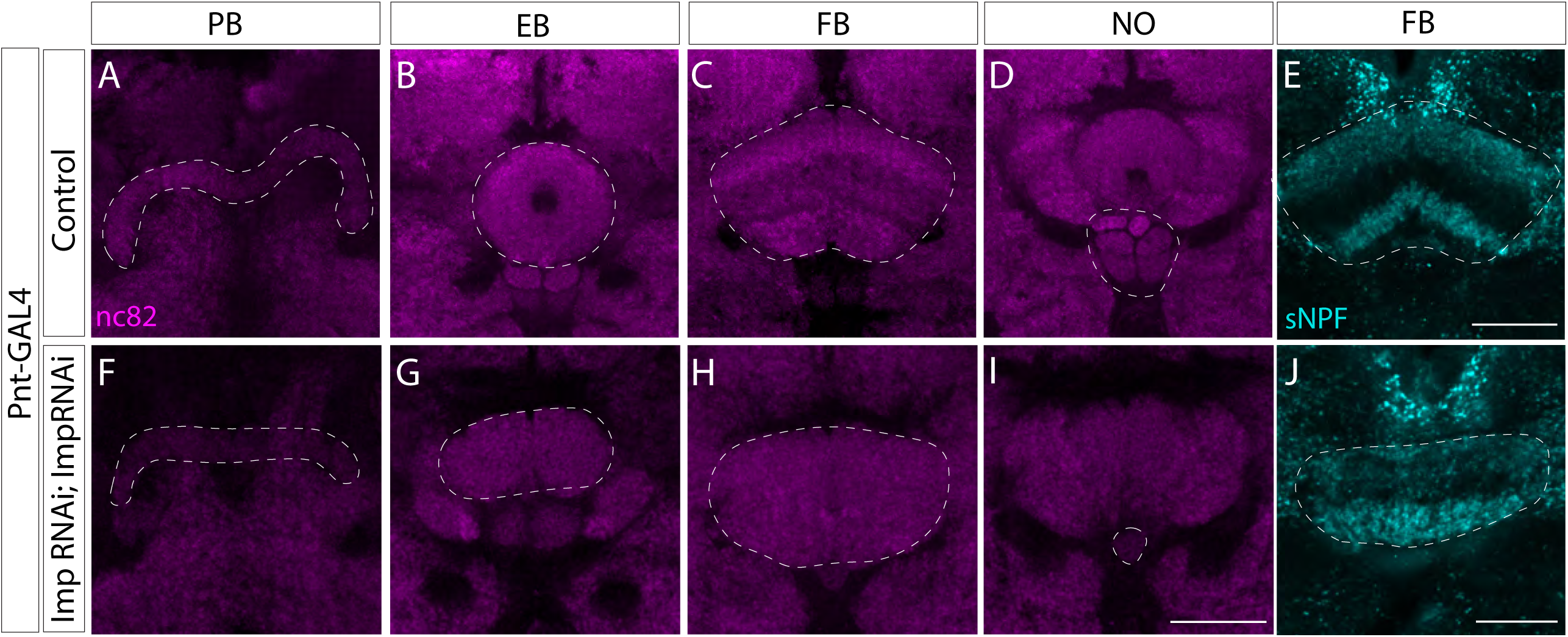
Imp regulates morphology of many neuropils and neuropeptide. **A-J**. The four neuropils of the CX: PB, NO, FB, and EB in control flies (A-D). Imp knock-down results in defective morphology of the CX neuropils **(F-I)**. nc82 stains the **E-J.** sNPF stains shown in cyan in control distributed in dorsal and ventral layers of FB distinctly **(E),** and Imp knock-down shows impaired distribution **(J)**. Scale bars correspond to 30μm.

Next, we wanted to examine the role of Imp in more broadly defining the neuropeptide distribution of FB layers. Previously we described expression of the neuropeptide TK (Fig. 4). We then examined the expression of short neuropeptide F (sNPF), which is reported to be strongly expressed in FB ventral layers 1-2 and dorsal layers 6-7^84^. In Imp knock-down flies, sNPF appears to be reduced in the dorsal layers (Fig. 6E and J). Together these results suggest that both the overall morphology of distinct neuropils as well as neuropeptide distribution of the CX are dependent on the levels of Imp in Type II neural stem cells.

### Imp is required in Type II neural stem cells for upwind orientation during olfactory navigation behavior

Given the profound effects of Type II-specific Imp expression on olfactory navigation circuitry, we sought to understand the impact of Imp expression on adult olfactory navigation behavior. We used Pointed-GAL4 to knock-down Imp in Type II neural stem cells, as in our anatomical experiments, using two copies of ImpRNAi. We then measured behavioral responses to a 10-second pulse of attractive odor (1% apple cider vinegar) using a previously described wind tunnel paradigm^79^.

Knocking down Imp in all Type II neural stem cells significantly impaired olfactory navigation behavior. In control flies (Pointed-GAL4>mCherryRNAi), odor evoked an increase in upwind velocity, an increase in groundspeed, and a decrease in angular velocity, while odor loss evoked an increase in turning (Fig. 7A, C), as observed previously^79^. Odor also increased the probability of movement, and this effect outlasted the stimulus by tens of seconds. In Imp knock-down flies (Fig. 7B, C), upwind velocity, ground speed, and angular velocity responses to odor were dramatically reduced. However, an odor-evoked increase in the probability of movement was still observed, arguing that the olfactory system is still functional and can influence motor behavior. To better understand these behavioral deficits, we decomposed upwind velocity during odor into two components: orientation upwind (Fig. 7D), and ground speed (Fig. 7E). Upwind orientation was strikingly decreased in Imp knock-down flies (Fig. 7D). In contrast, the distribution of groundspeeds during odor (for moving flies) was not substantially different for Imp knock-down flies versus controls (Fig. 7E). The trajectories of Imp knock-down flies also showed pronounced zig-zagging (Fig. 7B) highlighted by a greater average angular velocity (Fig. 7C) and a greater probability of making large amplitude turns (Fig. 7F). This suggests that in the absence of Imp, flies make more larger turns and have difficulty maintaining a stable heading over time.

**Figure 7.**
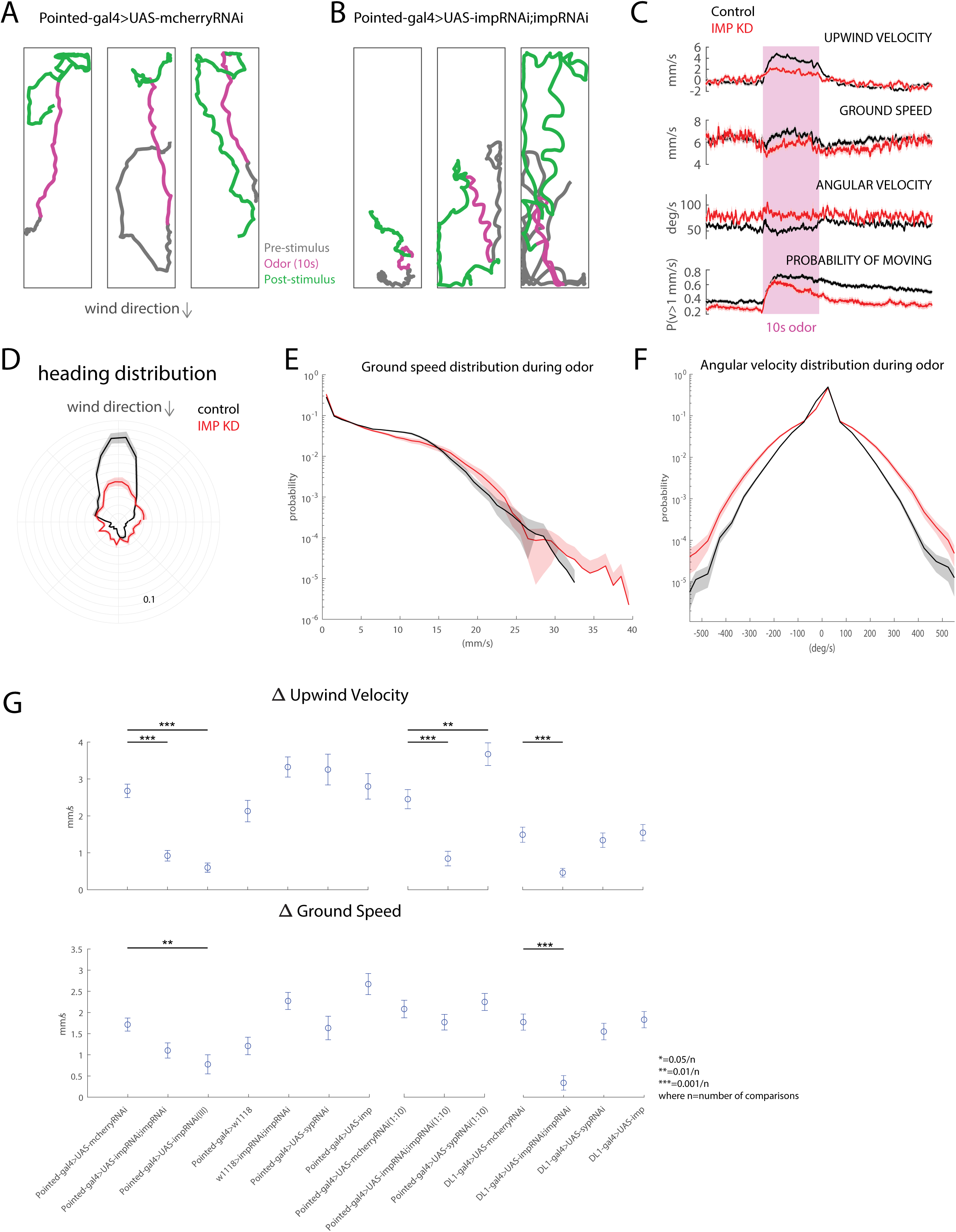
Imp regulates the upwind orientation during olfactory navigation. **A.** Representative walking trajectories of three different control flies (Pointed-gal4>UAS-mCherryRNAi) presented with a 10s odor pulse (1% ACV, magenta) centered in a 70s trial with consistent, 10cm/s wind. Flies demonstrate upwind movement during odor and local search after odor offset. Wind direction indicated above. **B.** Representative walking trajectories of three different Imp knockdown flies (Pointed-gal4>UAS-ImpRNAi;ImpRNAi) in the same wind and odor conditions as Figure A. Trajectories reveal increased turning behavior throughout the entirety of trial. **C.** Comparison of movement parameters between control (black, n = 86 flies) and Imp knockdown flies (red, n = 61 flies) calculated as an average across flies (mean±SEM; see Methods). Pink-shaded area: 10-second odor stimulation period (1% ACV). In the presence of odor, Imp knockdown flies show decreased upwind velocity comparable to control flies. Increased turning behavior noted in Figure B can be observed quantitatively through the increased angular velocity throughout the duration of trials. **D.** Heading distribution during odor for both control (black) and Imp knockdown (red) flies (wind direction indicated above). Imp knockdown flies exhibit a decreased probability of upwind orientation, corresponding to the decreased upwind velocity during odor stimulus seen in Figure C. **E.** Histogram showing ground speed distributions during odor for both control (black) and Imp knockdown (red). Distributions appear similar, suggesting that general movement is not impaired by Imp knockdown. **F.** Histogram showing angular velocity distributions for both control (black) and Imp knockdown flies(red) during odor. Imp knockdown flies favor larger angular velocities, even in the presence of an attractive stimulus. **G.** Change in upwind velocity and ground speed, calculated as the difference in the mean value during the first 5 seconds of odor and the 5 seconds preceding odor, across various control and experimental genotypes. Knockdown of Imp results in significant decreases in upwind velocity change when compared to respective controls, across multiple odor strengths and driver lines. Knockdown of late transcription factor Syncrip (Syp) also results in significant deficits in upwind velocity change. (two-sample t-test; from left to right, p = 5.906x10^-11, 1.2087x10^-8, 1.2293x10^-4, 0.0042, 9.0922x10^-6). Knockdown of Imp does not result in a significant difference in ground speed with the exception of DL1-gal4>UAS-ImpRNAi;ImpRNAi (two-sample t-test; p = 2.2281x10^-7). —------------------------------------------------------------------------------------------------------------------ - UPWIND VELOCITY pointed>mCherryRNAi compared to pointed>double ImpRNAi p = 5.906x10^-11 pointed>mCherryRNAi compared to pointed>ImpRNAi(III) p=1.2087x10^-8 pointed>mCherryRNAi (10:1) comp to pointed>double ImpRNAi (10:1) p=1.2293x10^-4 pointed>mCherryRNAi (10:1) comp to pointed>SypRNAi(10:1) p=0.0042 DL1>mCherry comp to DL1>double ImpRNAi p=9.0922x10^-6 GROUND SPEED DL1>mCherry comp to DL1>double ImpRNAi p=2.2281x10^-7

We observed similar effects on upwind velocity when using a single ImpRNAi transgene to perform the Imp knock-down (Fig. 7G), as well as when using a higher concentration of odor (10% vs 1% apple cider vinegar). We saw no change in upwind velocity modulation in parental strains (Pointed-GAL4 or ImpRNAi) crossed to wild-type. Type II stem cells are known to express opposing gradients of Imp and Syncrip over the course of development^77^. Knock-down of the late factor Syncrip produced no change in upwind velocity in response to 1% ACV but produced an increase in upwind velocity in response to 10% ACV. Finally, we examined the effect of knocking down Imp only in the DL1/DL2 Type II neural stem cells that generate tangential inputs to the CX. This manipulation also reduced upwind velocity (Fig. 7G). Upon deeper investigation, this effect appears to arise from lower overall ground speeds during odor presentation, as well as a weaker deficit in upwind orientation (Supp. Fig. 2). Overall, our behavioral investigations show that Imp knock-down significantly impairs olfactory navigation behavior while leaving basic locomotion (as measured by groundspeed distribution) and olfaction (as measured by the movement probability response) intact.

## Discussion

During development, neural stem cells generate diverse neuron types that assemble into distinct neural circuits enabling complex behaviors. Extensive work in genetic model systems has provided an overview of conserved temporal programs that govern the formation of diverse neuronal cell types ^65, 67, 86–88^. Although much is now known about the stem cell-specific molecular cues that determine cell type identity, understanding how these molecular cues shape the expression of complex behaviors is still in its infancy^89–91^. The insect CX provides an ideal model for dissecting the relationship between developmental processes and behavioral complexity. The majority of adult *Drosophila* CX neurons are derived from a few larval Type II neural stem cells, which follow a division program similar to that of cortical progenitors^59, 60, 63, 65–67^. The CX has been implicated in a number of behaviors, including olfactory navigation^45^, menotaxis^23, 43^, sleep^30–32^ and path integration in *Drosophila*^92–94^, and species-specific behaviors such as long-distance migration in monarch butterflies^95^, and allocentric dispersion in dung beetles^96^. The insect CX, therefore provides a powerful platform for understanding how changes to developmental programs give rise to changes in circuit structure and, thereby, behavior. We set out to study the developmental processes giving rise to olfactory navigation circuit elements and behavior in adult *Drosophila*.

### Lineage specific development of CX neuron types

Lineage based architecture plays an essential role in generating complex brain structures and circuits. In the CX, local and columnar neurons primarily arise from the dorsomedial lineages (DM1-4), while long-field tangential input neurons primarily derive from the dorsolateral lineage (DL1), with minor contributions from DM4 and DM6^53, 75^. Other major lineages that generate long-field tangential input neurons are the AOTU, LALv1, and SIPp1 which are Type I neural stem cells^53^. Our study reveals that major components of a previously described olfactory navigation circuit in the CX are all derived from Type II neural stem cells (Fig. 2). While many odor-encoding input neurons (vT029515) are born from DL1 neural stem cells, a few are derived from other DMs (Fig. 3). Our findings reveal that multiple lineages participate in building a complex circuit. *Drosophila* larval ventral nerve cord cohorts were shown to be formed by sequential addition from different lineages over time, where circuit input neurons are born prior to the output neurons^97^.

Our clonal analysis revealed that although tangential input neurons look morphologically similar, they are derived from different lineages; thus, there might be neuronal subtypes in that cluster, and each subtype might have distinct roles in generating a meaningful behavioral output. This also suggests that subpopulations in a neuronal class labelled by Gal4/LexA lines may be functionally heterogeneous. The DL1 lineage generates two distinct bundles, one innervating the dorsal and the other ventral layers of the FB. Based on our genetic birth-dating results, we found that the neurons that innervate the ventral layers and provide odor input to the olfactory navigation circuit are born between 48-72h (ALH) (Fig. 3). Our findings align with birth-dating studies on the entire DL1 lineage, which revealed that the ventral projecting neurons are born before 72h ALH, and the dorsal projecting neurons are born throughout larval development^77^. Whether the long-field tangential input neurons follow the conserved developmental principles across insects would be an interesting future study. Lineage analysis in grasshoppers and beetles has also shown remarkable similarities in the dorsomedial neural stem cells generating columnar neurons ^71, 73^. Whether other insects share a similar lineage-based architecture will be interesting to pursue. In the mammalian cerebral cortex, lineage is known to regulate neuron connectivity, where excitatory neurons originating from the same progenitors process related information and connect with each other, and inhibitory neurons are also known to organize in a lineage-dependent manner^98, 99^.

### Role of Imp in specifying an olfactory navigation circuit

Temporal gradients of two RBPs, Imp and Syncrip, have been shown to determine neural identity in the mushroom body, antennal lobe, and CX^77, 100^. In mushroom body neuroblasts, high Imp/low Syncrip levels early in development promote the specification of early-born y Kenyon cells, while low Imp/high Syncrip levels late in development promote the specification of late-born a/ý Kenyon cells^100–102^. In the antennal lobe, high Imp/low Syncrip leads to an increased number of late-born neurons from the antennal lobe anterodorsal 1 (ALad1) lineage at the expense of early born ones^100^. In the Type II lineages, DM1 and DL1 Imp/Syncrip levels control the number of several cell types^77^. However, the effects of these RBPs on multiple components of a functional circuit have not been previously characterized. Here we report a first step towards a developmental dissection of CX functional circuit formation and function. We found the strongest effects of Imp knock-down and overexpression on neurons targeting the ventral FB— ventral tangential inputs and ventral P-FNs. Curiously, Imp levels had distinct effects on each circuit element. Ventral FB tangential neuron numbers were increased or decreased through knock-down or overexpression of Imp, respectively (Fig. 4); in contrast, ventral P-FN numbers were reduced by Imp knock-down but not increased by overexpression (Fig. 5). The number of dorsal tangential inputs labeled by 65C03-LexA was decreased by both Imp knock-down and overexpression. These observations suggest that precise levels of Imp are required to specify the correct number of these neurons, and Imp might be acting in combination with other neural stem cell factors or INP factors to give rise to different cell types. Tangential input neurons targeting the dorsal layers of FB, FB5AB showed a decrease in cell body number upon Imp knock-down and no change upon Imp overexpression. Finally, the local FB neurons targeting the dorsal part of the fan-shaped body, hΔC/K did not change cell number with either Imp knock-down or overexpression, although their morphology was altered, suggesting altered function (Fig. 5). These results suggest that Imp plays multiple roles and acts via different molecular mechanisms in the neural stem cells to specify distinct neuron types from distinct lineages. In mice, Imp is expressed as a similar decreasing gradient in stem cells of CNS throughout embryonic development^103^, suggesting conserved mechanism. Imp has been previously reported to play a role in stem cell proliferation and tumorigenesis^82, 104–106^, however we report distinct effects of Imp on multiple circuit elements ruling out a generalized phenomenon. In the ventral nerve cord motor neuron lineages, opposing gradients of Imp and Syncrip in post-mitotic neurons regulate precise axonal targeting^107^. We here show that the expression of Imp post mitotically in vT029515 neurons maintains their identity (Fig. 4E-F). Fate specification and maintenance in post mitotic neurons has been previously reported in *C. elegans*^87, 108^ and *Drosophila* optic lobe^109^ and motor neurons as well^110^.

Many neuropeptides express in the CX in distinct patterns and modulate various behaviors, including locomotion, feeding, and olfaction^84, 111–115^. Here, we show that ventral FB tangential input neurons labeled by VT029515 express the neuropeptide TK and that Imp expression in Type II neural stem cells is necessary for regulating the neuropeptide identity of these neurons (Fig. 4). Moreover, we find that manipulating Imp levels in Type II neural stem cells affects the morphology of CX neuropils and distributions of two neuropeptides, TK and sNPF, throughout the FB (Fig. 6). These studies provide the first insights into the relationship between developmental timing and establishing neuropeptidergic identity in CX. Whether other temporally expressed developmental factors regulate the neuropeptide identity of CX lineages remains to be tested. In mice, loss of Imp leads to deformities in the posterior brain, neuroepithelial orientation defect, and cellular packing deficiency with poorly defined barriers between cortical layers/zones^85^, similar to the lamination defect we observe in FB and overall defective morphology of CX.

One open question is how Imp regulates neural fate and the function of multiple circuit elements at the molecular level. RBPs are known to play essential roles in regulating temporal gene expression by affecting the stability and translation of mRNA^100^. In mushroom body and antennal lobe stem cells, Imp regulates cell fate by post-transcriptional regulation of the transcription factor Chinmo^100^. Previous studies have shown that Chinmo is persistently expressed in Type II neural stem cells that are mutant for Syncrip and thus maintain high Imp expression throughout development, and this manipulation resulted in the formation of extra early-born neuron types^76, 77^. In the ventral cord motor neuron lineage, LinA/15, Imp, and Syncrip regulate axon-muscle connectivity by regulating various transcription factors post-transcriptionally^107^. Although precocious expression of Imp in Type II neural stem cells generated extra VT029515 neurons, to our surprise, we didn’t observe any changes in VT029515 neuron numbers upon Syncrip knock-down (Supp. Fig. 3). In addition, Type II neural stem cells generate INPs that inherit Imp, and the transcription factors expressed sequentially in INPs might act in combination with Imp to specify distinct neural fates. Type II neural stem cell transcription factors might also regulate the INP temporal transcription factor code. Future studies examining the downstream targets of Imp will reveal the exact mechanisms underlying Imp’s effects on cell number and morphology. Previous work has also shown a role for Imp in the timing of neural stem cell quiescence and proliferation^83, 106^. Since many other neural cell types that arise from Type II neural stem cells are not affected upon Imp knock-down, we conclude that the phenotypes we observe are less likely to be associated with quiescence or cell proliferation. RBPs can regulate gene expression by creating liquid phase separation granules. Imp has intrinsically disordered domains and is known to promote phase separation in mushroom body neurons^116^. Whether Imp makes phase-separated granules in Type II neural stem cells and if that has a role in fate specification is not known. Future studies will elaborate on the different molecular mechanisms underlying the effect of Imp on multiple circuit elements.

### Developmental dissection of CX-mediated behavior

In this study, we show that knock-down of the early-expressed RBP Imp in Type II neural stem cell lineages largely abolish the upwind orientation in response to odor. These flies show a marked inability to maintain a straight orientation, with larger turns and "zig-zagging" behavior both at baseline and in the presence of odor. This phenotype is reminiscent of classical CX mutants such as no-bridge-KS49 and CX-KS181^8^, which likewise showed an inability to maintain locomotor fixation, in this case towards a visual target. Intriguingly, we did not observe any deficit in walking velocity in our knock-down flies, suggesting that basic locomotion is intact. Imp knock-down flies also showed increased locomotion in response to the odor pulse, suggesting that olfactory function is intact. Knock-down of the late-expressed RBP Syncrip did not impair olfactory navigation behavior and even increased upwind velocity in certain conditions, arguing that neural circuits established early in CX development are critical for this behavior. Taken together, our results suggest that CX circuitry established early in development is critical for maintaining a consistent heading, in particular towards a wind cue, but potentially to other cues as well (Fig. 7).

Current approaches to CX circuit dissection in fruit flies have emphasized the use of highly-specific GAL4 and Split-GAL4 lines, which target specific subsets of neurons^117, 118^. Silencing of highly specific CX lines often produces fairly subtle phenotypes. For example, silencing of compass neurons eliminates menotaxis (orientation to a visual stimulus at an angular offset) but not visual fixation per se^23, 119^, while silencing of hΔC/K neurons impairs the persistence of upwind tracking behavior during odor, but not the initial upwind turn^45^. As neurons with related functions may arise from the same progenitor, stem-cell based manipulations that target developmentally related neurons may produce more striking phenotypes and provide complementary insight into the functional organization of neural circuits. Future experiments manipulating other temporally expressed transcription factors or intersecting their manipulation with the temporal transcription factor code in INPs, should allow us to make more precise manipulations of CX development.

Here we showed that Imp knock-down in Type II stem cells produces diverse changes to CX organization. At a macro level, several neuropils show altered morphology, and the expression of peptides in particular layers of the FB is disrupted. At a micro-level, inputs to the ventral FB that normally carry wind direction and odor information are absent, while dorsal FB neurons thought to carry a "goal" signal for olfactory navigation show impaired morphology. A working hypothesis is that the behavioral deficits we observed might arise from the loss of wind and odor sensory inputs to the ventral FB with additional contributions from morphological deficits of other circuit components. In addition, disruption of the global heading signal in the EB^15^, or the loss of peptide signals in the FB^114^ could contribute to the observed behavioral deficit. Mushroom body output neurons that innervate the vertical lobes (MBONs 15-19)^120^ and drive upwind orientation^45^ also arise from the DL1 neural stem cell^121^, and could play a role in the phenotype we observed with Imp knock-down. Future experiments examining neural encoding and optogenetic activation in developmentally altered circuits will provide insight into the precise circuit basis of this behavioral deficit.

## Acknowledgments

The authors would like to thank Chris Doe and Doe lab members for their valuable feedback and discussions. We thank Chris Doe, Raees Andrabi, and Dena Goldblatt for providing critical feedback on the manuscript. We are grateful to Chris Doe, Claude Desplan, Gerry Rubin and Tzumin Lee for sharing the reagents. Stocks obtained from the Bloomington Drosophila Stock Center (NIH P40OD018537) were used in this study. The monoclonal nc82 antibody was obtained from the Developmental Studies Hybridoma Bank, created by the NICHD of the NIH and maintained at The University of Iowa, Department of Biology, Iowa City, IA 52242. We thank UNM Biology cell biology core for providing confocal microscopy facility. N.C and A.G were supported by NIH URISE T34 GM145428. The research was supported by NSF 2014217, NIH R01DC017979 and RF1NS127129 to K.I.N and National Science Foundation CAREER Award IOS-2047020 to MHS.

## Author contributions

Conceptualization, A.H, K.I.N, and M.H.S.; methodology, A.H performed all anatomy experiments and confocal imaging with help from A.G, H.G, A.N.C, M.P, and T.S. performed behavior experiments; visualization, A.H, K.I.N and M.H.S.; writing, A.H.; editing, K.I.N, and M.H.S; funding acquisition, K.I.G, and M.H.S; and supervision, K.I.N, and M.H.S.

---

**Figure Supplement 1.**
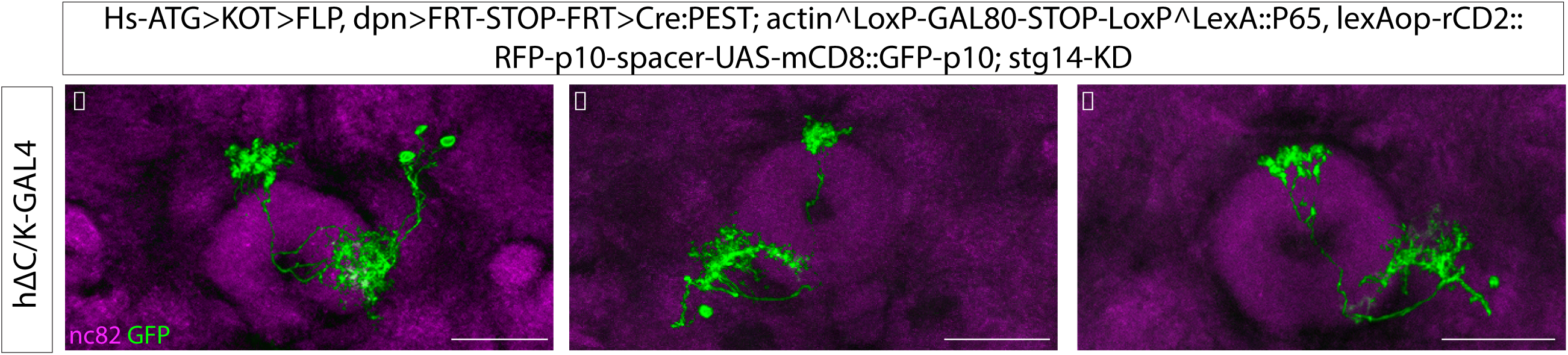
CLIn clones of hΔC/K neurons. **A-C**. Confocal images of CLIn crossed to vT062617 GAL4 shows this driver line labels hΔC/K neurons that innervate FB and EB. hΔC/K neurons are labelled in green (GFP), nc82 counterstains are used to visualize EB. The genotypes are shown at the top and the abbreviated genotypes for neuron class is shown at left. Scale bars correspond to 30μm.

**Figure Supplement 2.**
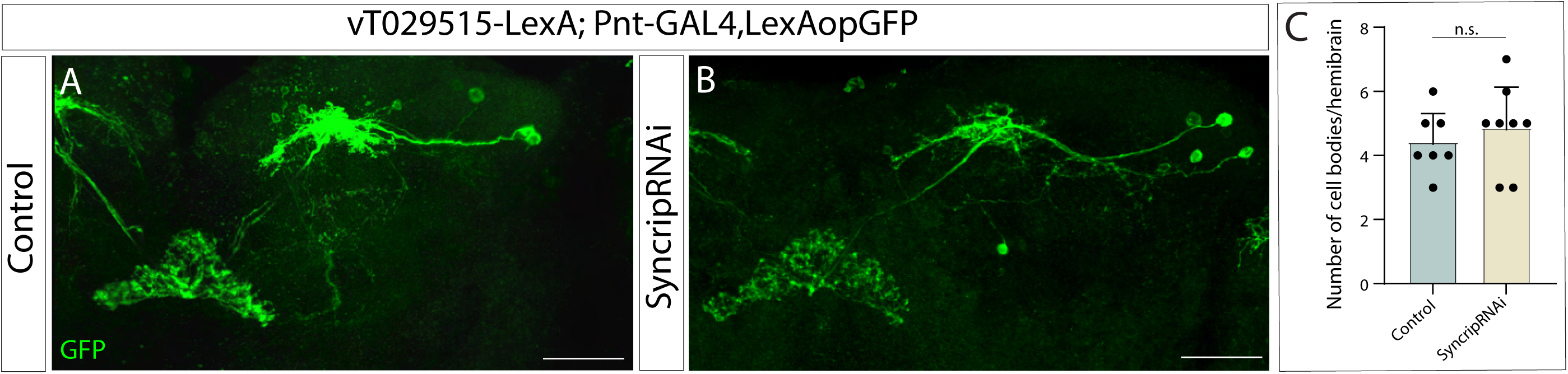
Knock-down of Syncrip does not change the number of vT029515 neurons. **A-B**. vT029515 neurons labelled by reporter GFP in control **(A)**, and Syncrip knock-down **(B)**. GFP is shown in green. The abbreviated genotypes are shown at the left and top, respectively.

**Figure Supplement 3.**
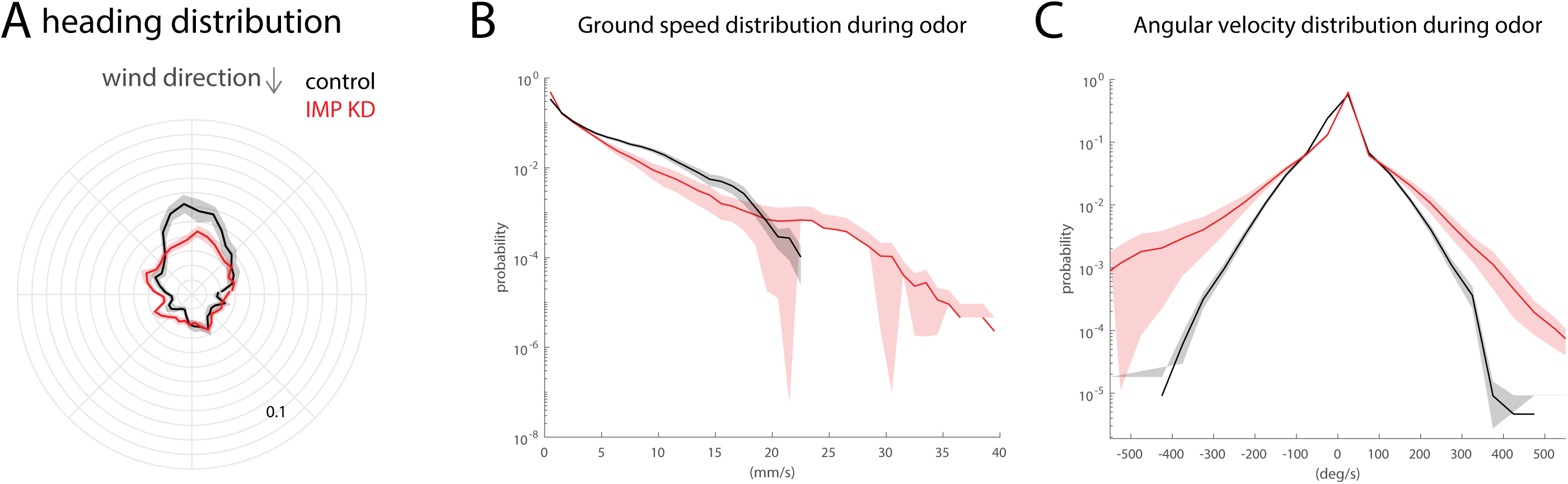
Knock-down of Imp in DL1/2 neurons impacts both orienting ability and speed modulation. **A.** Heading distribution during odor for both control (black) and knockdown of Imp (red) in DL1/2 neurons (wind direction indicated by above arrow). **B.** Histogram showing ground speed distribution during odor for both control (black) and Imp KD (red) flies. Imp KD flies tend towards lower ground speeds than control counterparts. **C.** Histogram showing angular velocity distribution during odor for both control (black) and Imp KD (red) flies. Widening of distribution indicates that Imp KD flies favor extraneous angular velocities.

## Materials and methods

### Experimental model details

For all experiments, we used *Drosophila melanogaster*. All the fly stocks were maintained at 25°C on standard cornmeal-agar medium and a 12h light-dark cycle. The RNAi and overexpression experimental crosses were raised at 29°C. Adult flies dissected were 3-7 days old. For behavioral experiments, female flies were collected at least 1 day after eclosion and then placed in vials that were situated in custom-made cardboard boxes at room temperature, with a 12-hour light-dark cycle, for at least two days to acclimate to ambient temperature and ensure correct circadian rhythm for the experimental time of day. All experiments were conducted within 4 hours after the flies’ apparent “dawn” (ZT 1-ZT 4). The flies were then starved for 24 hr in an empty transparent polystyrene vial that contained a Kimwipe that was soaked in distilled water to humidify the air. By the day of the behavioral experiments, the flies had an average age of 3 to 7 days. Flies were anesthetized over ice for approximately 1 minute before being loaded into wind tunnels and allowed at least 5 minutes to recover before starting the first trial. The exact genotypes for each figure panel are listed in the table below. Parental strains and RRIDs are listed in the Key resources table.

### Immunostaining

Adult fly brains were dissected in Schneider’s Insect medium (Sigma-Aldrich), and then fixed at room temperature for 23 min in 4% paraformaldehyde (EMS) in PBT buffer. 10X PBS buffer stock contained NaCl: 1.37 M, KCl: 27 mM, Na2HPO4: 100 mM, KH2PO4: 18 mM with a pH of 7.4. Working 1X PBT had an addition of 0.5% TritonX-100 (Sigma-Aldrich). Fixing was followed by washing at room temperature in PBT to remove PFA. The samples were blocked at room temperature for 40 min in PBT containing 2.5% normal goat serum (Jackson ImmunoResearch) and 2.5% normal donkey serum (Jackson ImmunoResearch). Adult brains were then incubated for 48hr at 4°C in the primary antibody solution.

Primary antibody staining was followed by washing in PBS-T and then the adult brains were incubated for 2hr at room temperature in the secondary antibody solution. The brains were then washed in PBT. The primary and secondary antibody solutions were prepared in the blocking solution (prep described above).

Following the antibody staining, brains were mounted on the Poly-lysine-coated cover slips, serially dehydrated in alcohol with concentrations 30%, 50%, 75%, 95% and 3 rounds of 100% to replace PBT. The brains were then cleared in Xylene with 3 rounds of 5 min each and then mounted in DPX mounting medium, dibutyl phthalate in xylene (Sigma-Aldrich #06522). The samples were allowed to set in DPX for 3 days before imaging.

### Microscopy

Images were acquired using a Zeiss LSM780 confocal microscope, analyzed using ImageJ, and processed using ImageJ and Photoshop. Cell body numbers were calculated manually using a cell counter in ImageJ, and the statistical analysis and graphs were made using GraphPad Prism. Two-tailed student t-tests were used to compare control with loss of function and gain of function respectively for cell body numbers. Asterisks denote levels of significant differences *p<0.05; **p<0.01; ***p<0.001; ****p<0.0001.

### Birth-dating using Cell class lineage intersection system

Males from neuron class specific GAL4 lines were mated to females from the CLIn line^81^. Egg collection was done for 3 to 4 hr intervals on apple juice containing agar plates. Hatched larvae (0-3 hr old) were manually collected and raised in standard fly food at 25°C. Heat shock was applied to the hatched larvae for 12-15 min for lineage analysis and 40 min for birth-dating at the time point 0 hr, 24 hr, 48 hr and 72 hr ALH. F1 adult flies of age 3-7 days were dissected.

### Behavioral Apparatus

All behavioral experiments were performed in miniature wind tunnels^79^. Flies were constrained to walk in a shallow acrylic arena with a constant laminar wind at 10 cm/second. The wind tunnels were backlit using IR LEDS (850 nm, Environmental Lights) and movement was recorded using a camera placed below the chamber (Basler acA1920-155um). A 10 s pulse of odor (either 1% or 10% apple cider vinegar) was presented through Lee valves connected to the wind tunnel with polyethylene tubing. Within each 70 s trial, flies experienced 30 s of clean wind, 10 s of odor with wind, and then 30 s of clean wind again. All stimuli were controlled through a NIDAQ board. There are roughly 5 seconds between the end of one trial and the start of the next. Position (x,y) and orientation data were computed and collected in real time using a double-thresholding algorithm as described previously using NI LabVIEW).

### Analysis of Behavioral Data

All analysis was completed using MATLAB (Mathworks, Natick, MA). Calculation of trajectory and basic movement parameters seen in Fig. 7. A-C (ground speed, orientation, etc.) were completed using analysis code described in Alvarez-Salvado *et al*, 2018. Trials were excluded as a result of tracking error or if flies did not move a minimum distance of 25mm throughout the duration of the trial.

### Orientation histograms

To compute orientation histograms, we pooled trials across all flies, and separated orientation data from the period before odor presentation, the period during the 10 seconds of odor, and the period following odor offset. We binned orientation data from the first 5 seconds of each of these periods into 36 bins of width 10 degrees. Error bars were created using jackknife resampling across flies and represent standard error calculated as:

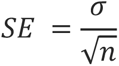

where *σ* is standard deviation and *n* is the number of samples. The resultant data, in polar coordinates, was then converted to Cartesian coordinates for ease of plotting and creation of error bars.

### Histograms of ground speed and angular velocity

To compute groundspeed and angular velocity histograms, we again pooled data across flies. We binned values of ground speed and angular velocity into bins of width 1mm/s and 50 deg/s respectively (Fig. 1. E, F) Histograms of both parameters were normalized to probability. Error bars represent standard error and were created using jackknife resampling with bias correction across all flies. Angular velocity of stationary flies (ground speed< 1mm/s) were excluded from analysis.

### Summary plots of change in forward velocity and ground speed

Iterating through all trials for a single fly, we calculated the difference between the mean upwind velocity in the first 5 seconds of odor presentation and the mean upwind velocity in the 5 seconds preceding odor presentation. We then calculated the mean change in upwind velocity between these two periods for a single fly by averaging across all trials. For data representation, we then calculated the mean value across all flies, as well as the standard error. Significance was calculated using the mean value of each fly as a data point in an unpaired t test with Bonferroni correction for multiple comparisons using the equations:

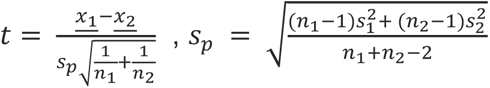

where <u>*x*_1_</u> and <u>*x*_2_</u> are sample means, *n*_1_ and *n*_2_*n*, are sample sizes, *s*_p_ is the population variance, and *s*_1_ and *s*_2_ are the sample variances. The same logic was used for calculating change in ground speed and performing statistical testing (Fig. 1 G). Significance is marked with asterisks in the figure and the associated p-values can be found below.

### Upwind Velocity

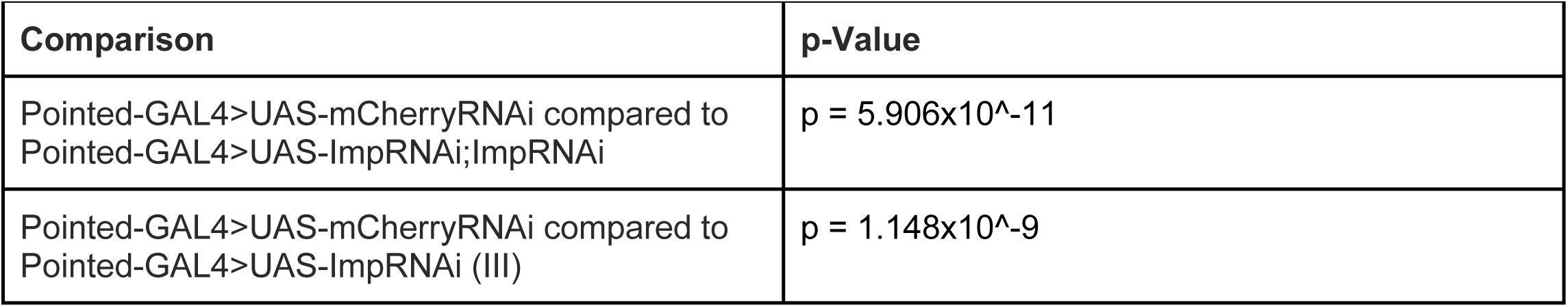
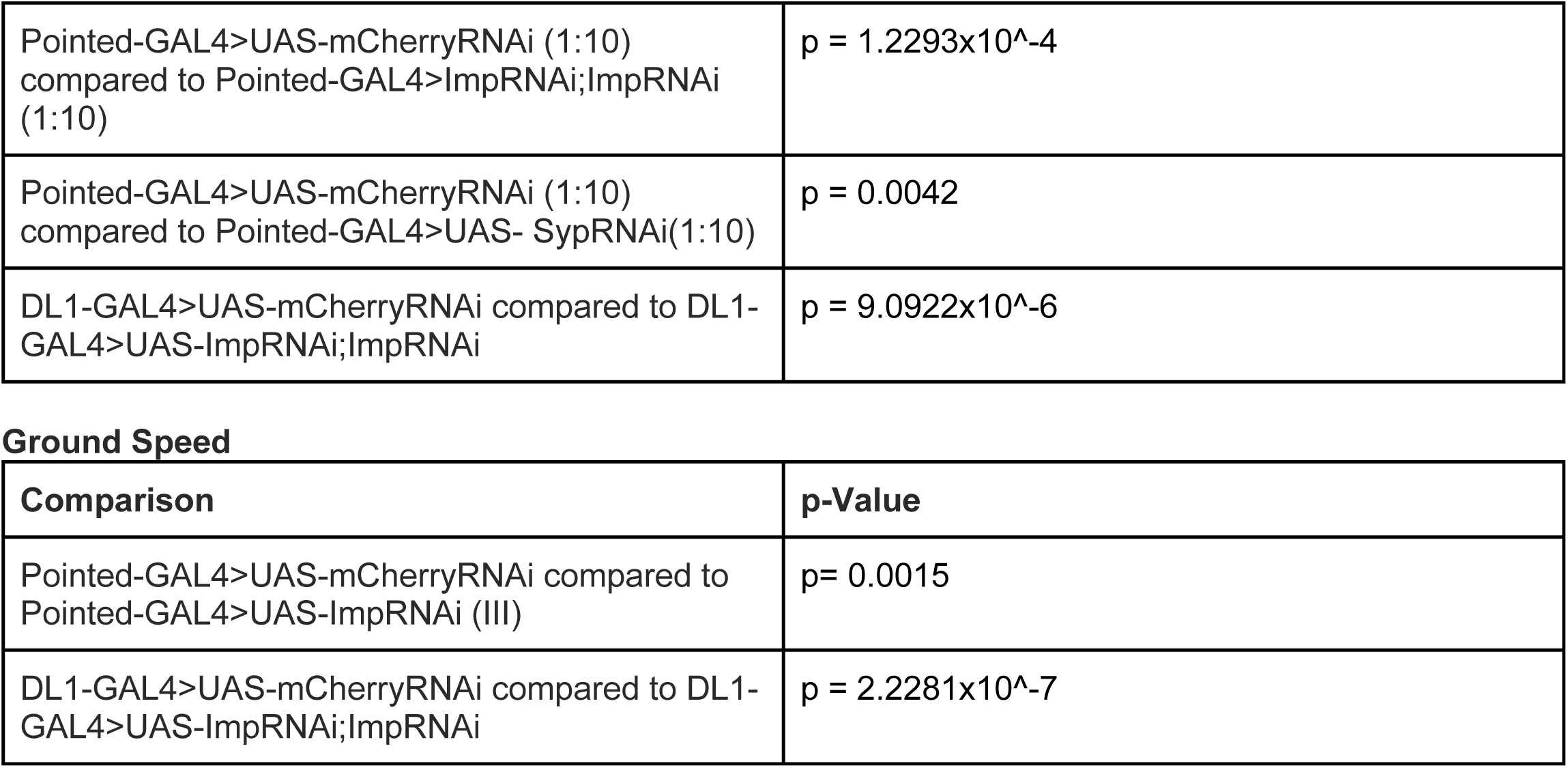

| Comparison | p-Value |
| --- | --- |
| Pointed-GAL4>UAS-mCherryRNAi compared to Pointed-GAL4>UAS-ImpRNAi;ImpRNAi | p = 5.906x10<sup>-11</sup> |
| Pointed-GAL4>UAS-mCherryRNAi compared to Pointed-GAL4>UAS-ImpRNAi (III) | p = 1.148x10<sup>-9</sup> |
| Pointed-GAL4>UAS-mCherryRNAi (1:10) compared to Pointed-GAL4>ImpRNAi;ImpRNAi (1:10) | p = 1.2293x10<sup>-4</sup> |
| Pointed-GAL4>UAS-mCherryRNAi (1:10) compared to Pointed-GAL4>UAS- SypRNAi(1:10) | p = 0.0042 |
| DL1-GAL4>UAS-mCherryRNAi compared to DL1-GAL4>UAS-ImpRNAi;ImpRNAi | p = 9.0922x10<sup>-6</sup> |

Ground Speed
| Comparison | p-Value |
| --- | --- |
| Pointed-GAL4>UAS-mCherryRNAi compared to Pointed-GAL4>UAS-ImpRNAi (III) | p= 0.0015 |
| DL1-GAL4>UAS-mCherryRNAi compared to DL1-GAL4>UAS-ImpRNAi;ImpRNAi | p = 2.2281x10<sup>-7</sup> |

### Fly Strains

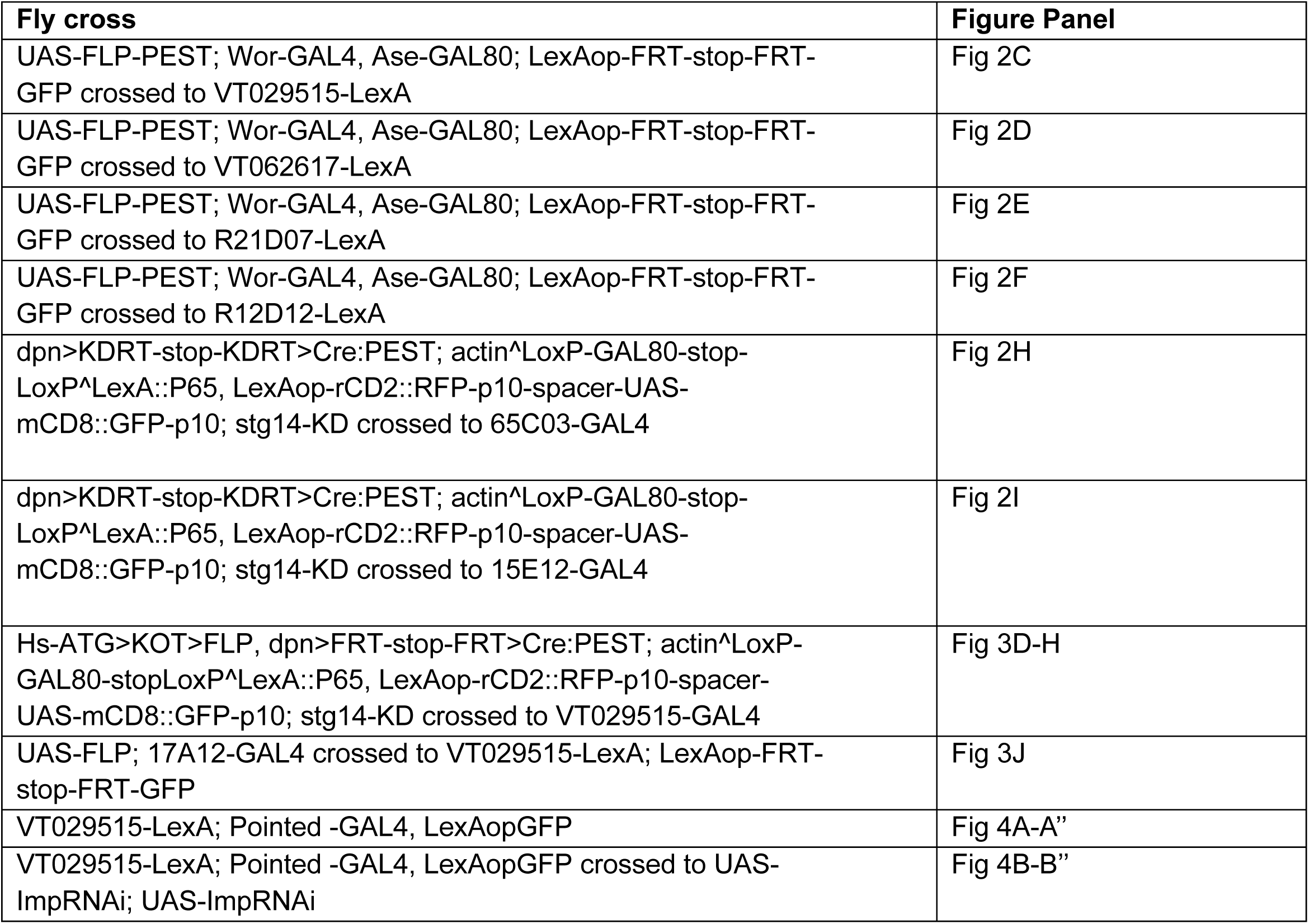
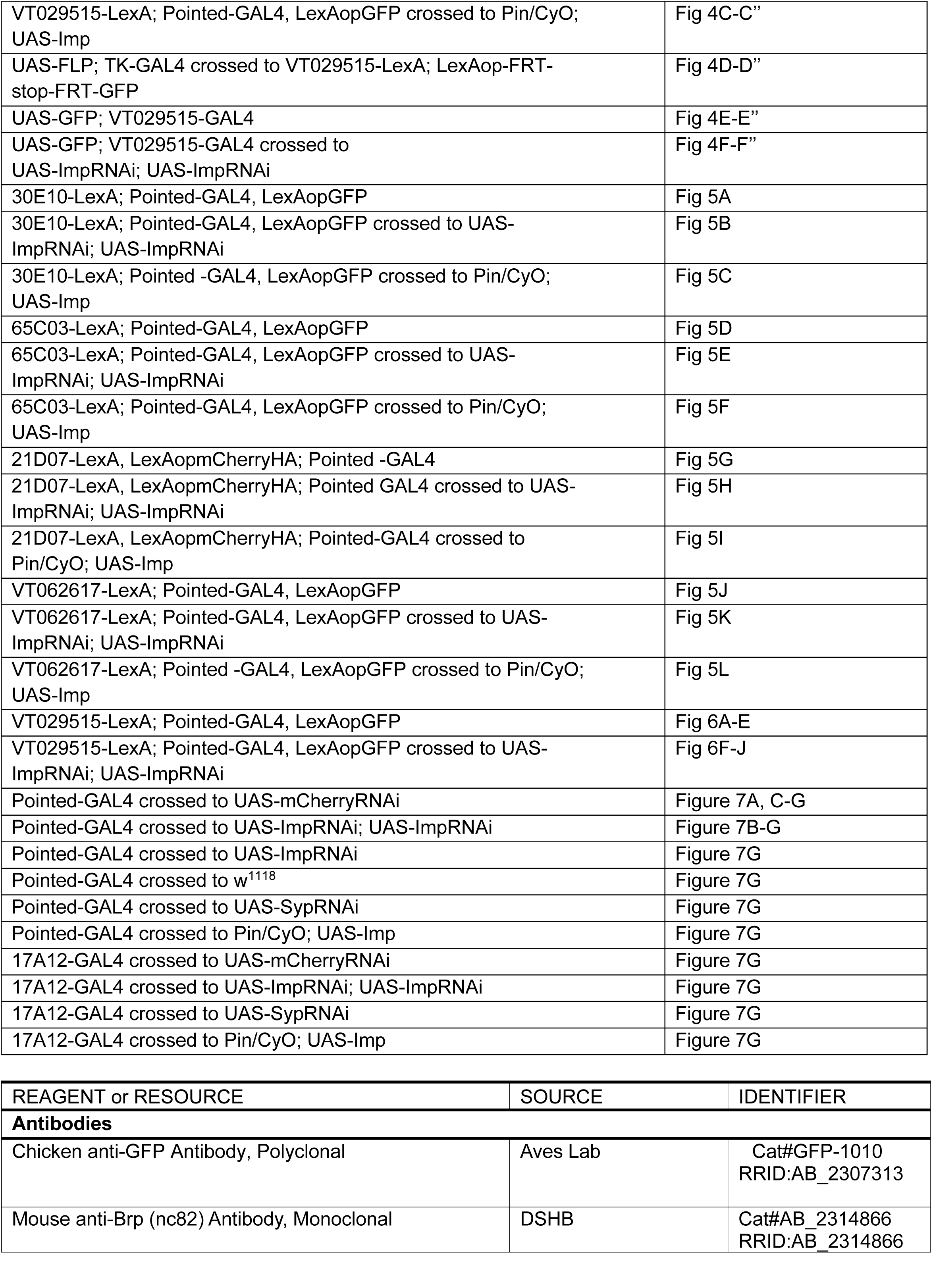
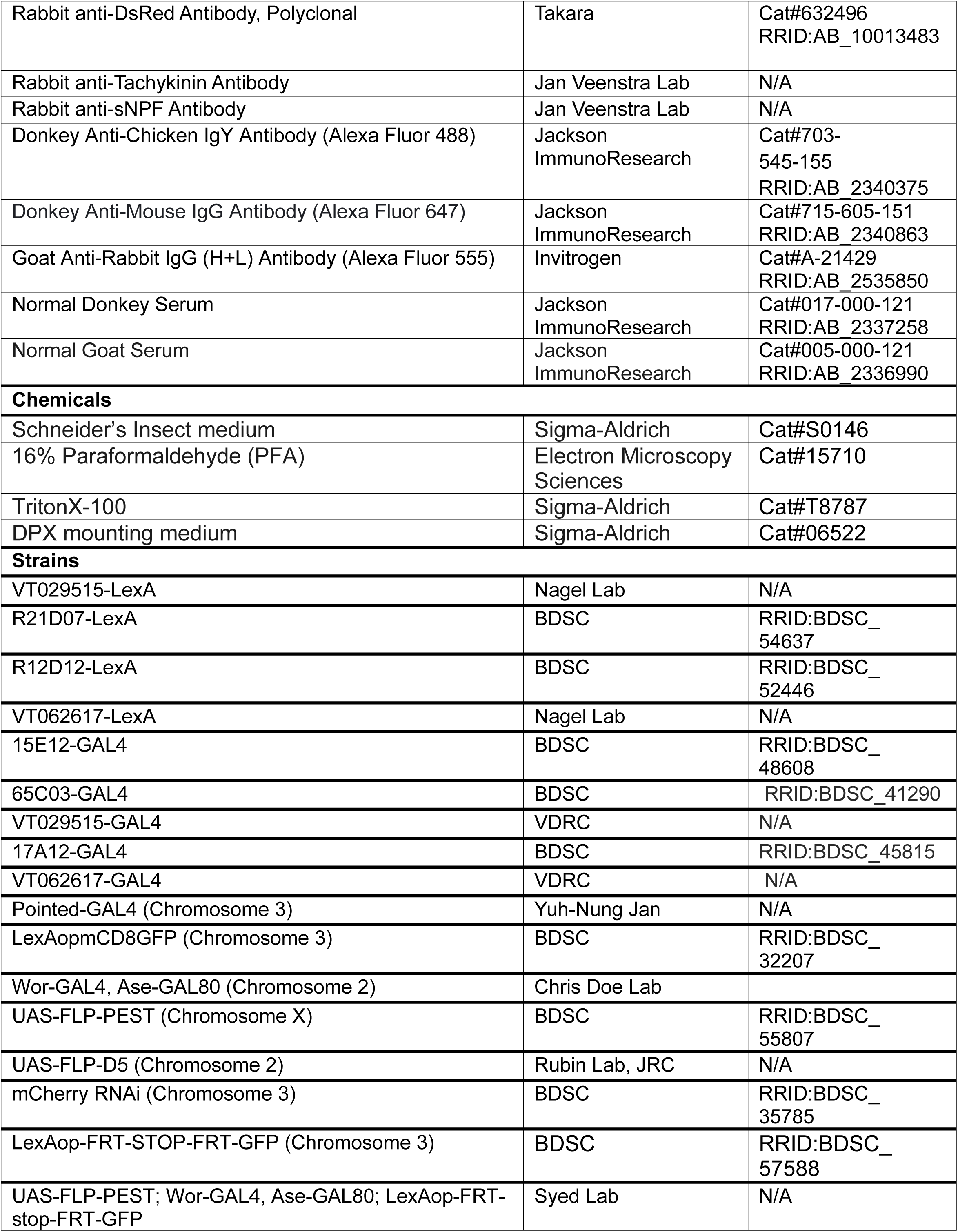
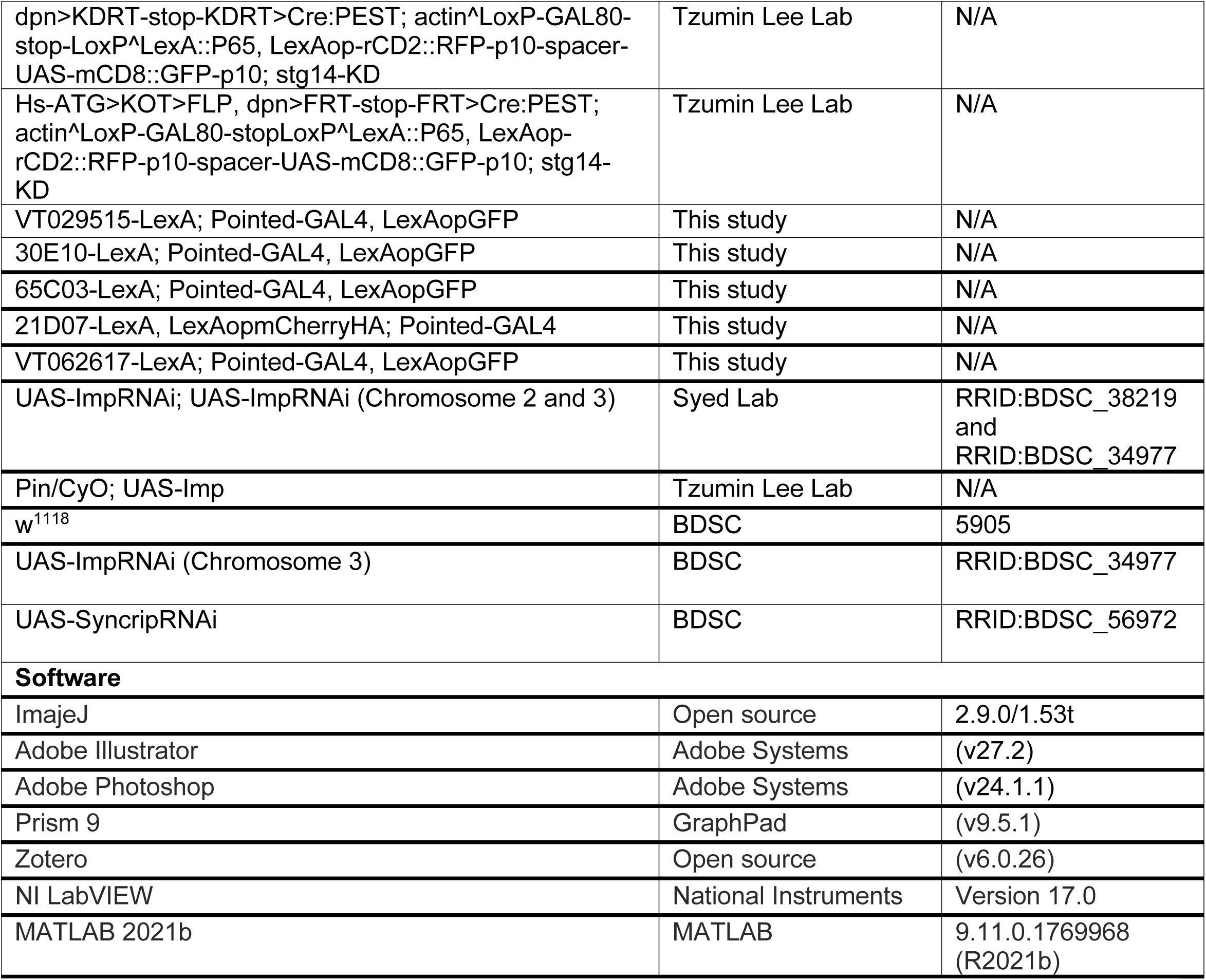

